# Urban planning leaves an audible legacy in a megacity’s bird soundscape

**DOI:** 10.64898/2026.10.18.759919

**Authors:** Yuhao Chen, Zezhou Hao, Chengyun Zhang, Zongying Li, Lyubing Zhang, Zihao Chen, Wenqiang Fang, Wei Liang, Ting Zhou, Yang Liu

## Abstract

Cities are planned for people, yet planning also determines where wildlife persists and where residents hear it. How can this acoustic footprint guide planning and monitoring? We combined AI-powered passive acoustic monitoring at 31 stations with detection-corrected models and spatial validation to map Asian Koel activity across Shenzhen. Koel acoustic opportunity declined with tree and built cover and increased with starling and myna opportunity; host information improved prediction at unmonitored sites. Weekly opportunity expanded in spring and varied across the city; sampled-minute exposure concentrated around dawn. Supported predictions intersected planning-sensitive places, while 51.5% of the city grid remained a monitoring gap. We translated these boundaries into monitoring priorities for underrepresented environments and uncertain spring predictions. This reproducible workflow reveals the acoustic imprint of the present-day urban landscape and connects ecological diagnosis, prediction boundaries and adaptive monitoring to inform urban coexistence.

## Introduction

Urban planning shapes where wildlife persists—and where people hear it (*1*, *2*). As cities expand, decisions about parks, vegetation, buildings and infrastructure reorganize habitats and bring wildlife into daily contact with dense human populations (*1*, *3*). Over time, these choices assemble an audible city: some animal sounds enrich everyday encounters with nature, while conspicuous seasonal calls can become recurring sources of tension (*4–6*).

One example is the Asian Koel (*Eudynamys scolopaceus*) in Shenzhen, a megacity shaped by four decades of exceptionally rapid urbanization. Now Shenzhen confronts this consequence through the Asian Koel, whose powerful breeding-season calls have prompted resident complaints, local-authority responses and repeated media attention (7–9; Fig. 1B). This controversy exposes two blind spots in urban planning. First, cities create landscapes that support wildlife, yet rarely know how habitat configuration, ecological interactions and seasonal behavior combine to place a particular sound near homes, parks, schools or hospitals (*1*, *10*, *11*). Second, sparse monitoring leaves planners uncertain about where citywide predictions are reliable and where new observations are needed (*12*, *13*). Closing these gaps requires two urgent tasks: diagnosing how the present-day landscape, shaped over decades of rapid urban development, organises acoustic activity, and using that diagnosis to design future urban spaces together with the monitoring network needed to evaluate them.

**Fig. 1.**
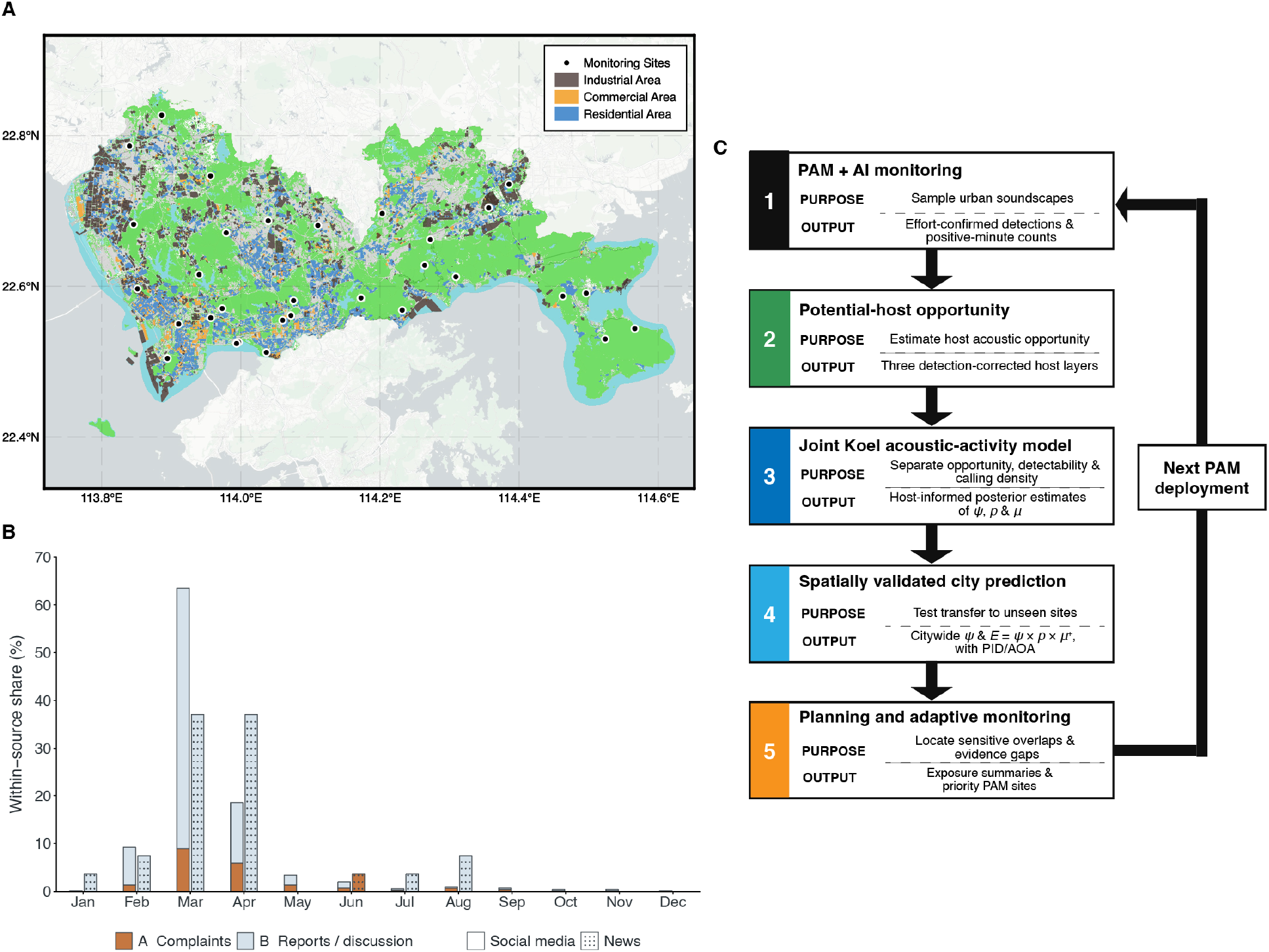
Study area, public attention and analytical workflow for citywide Asian Koel acoustic monitoring. (A) Locations of the 31 passive acoustic monitoring (PAM) stations across Shenzhen. Black circles mark stations; brown, orange and blue polygons indicate industrial, commercial and residential areas, respectively. (B) Calendar-month distribution of nuisance-related social-media posts and news records published from January 2023 to September 2026. Social-media records (n = 712) and news records (n = 27) are summarised separately. Orange denotes personal complaints or self-reported disturbance (A), and pale blue denotes relevant reporting, retelling or discussion (B). Unpatterned bars indicate social media and dotted bars indicate news. For each group, monthly A and B counts were divided by its total dated A+B records; both categories and all months together sum to 100%. Social-media counts were pooled before calculating percentages. (C) Workflow linking AI-labelled PAM recordings, detection-corrected potential-host opportunity, joint Koel acoustic-activity modelling, spatially validated citywide prediction and adaptive monitoring. The joint model separates weekly acoustic opportunity (*ψ*), daily detectability (*p*) and calling density (*μ*); sampled-minute acoustic exposure is *E* = *ψ* × *p* × *μ*^+^, where μ⁺ is the zero-truncation-adjusted positive-day sampled-minute probability. Prediction Interpretation Domain (PID) and Area of Applicability (AOA) labels identify prediction support, while planning-sensitive overlaps and evidence gaps guide subsequent deployment. Media screening and aggregation are described in Supplementary Methods S1.4; station and spatial-layer information is provided in Supplementary Data 1, 7 and 15.

The Asian Koel brings these two tasks into focus: its dependence on suitable hosts and conspicuous seasonal calls offer a clear case for linking landscape diagnosis with future planning and targeted monitoring. As an obligate brood parasite, it must find nests of suitable hosts, and experiments show that Koels can locate host nests even at unfamiliar sites (*14*). The identity of those hosts shifts across Asia: records include mynas and crows on the Indian subcontinent, multiple species—predominantly Corvids—in China, and chiefly House Crows (*Corvus splendens*) in Singapore (*15–19*). Host availability therefore defines a moving ecological opportunity that urban landscapes can concentrate or disperse. Male Koels deliver loud, distinctive calls during the breeding season, with pronounced activity around dawn and dusk, making their breeding activity a highly audible event of urban life (*16*). Public responses are divided: the same calls are recognised as part of urban nature and reported as a recurrent disturbance in Asian cities (*8*, *15*, *20*). The Koel’s ecology identifies three key factors—hosts, habitat and season (*14*, *16*). These factors raise a citywide question: why does acoustic activity concentrate in particular places and seasons, and can information of habitat and potential-hosts improve prediction beyond landscape and season alone? Citywide evidence must then show where that activity intersects planning-sensitive places and whether patterns learned from monitored neighbourhoods hold elsewhere (*10*, *21*).

Passive acoustic monitoring (PAM), coupled with AI, can build that evidence at the necessary temporal scale (*22–24*). Autonomous recorders revisit the same places across seasons and the daily cycle, while machine-learning systems turn sound archives into repeatable species observations (*25*, *26*). This combination has already enabled urban ecoacoustic assessment, detection-corrected population inference and mapping of soundscapes and vocal activity (*11*, *27*, *28*). Yet a recorder observes a point, while a planner must act across a surface (*29*, *30*). Scaling transfer from monitoring stations to citywide maps therefore requires three safeguards. First, models must separate weeks without Koel activity from weeks in which Koels were active but went undetected (*31*, *32*). Second, transfer must be tested at stations withheld from model fitting, because random folds can reward spatial memorisation and overstate performance (*21*). Third, every prediction must be checked against the environments represented by the monitoring network; applicability analysis identifies supported locations and flags environmental extrapolation (*12*). Together, these safeguards turn a visual extrapolation into evidence with a known domain. Planning imposes a further demand: the map should identify the observations that would make its next version stronger. Adaptive monitoring formalises this learning cycle by connecting evidence, uncertainty and the next sampling decision (*33*, *34*). PAM becomes significant urban infrastructure when it can map where wildlife is acoustically active, state where that map deserves confidence and direct the next recorders towards the city’s most consequential evidence gaps.

Evidence is useful for planning when it connects ecological activity with human experience (*6*, *10*). Bird sounds can support restoration, well-being and connection with nature, but their effects depend strongly on species, place and time (*4*, *35*, *36*). Conspicuous calls can create recurring coexistence pressure where seasonal activity overlaps residential areas, schools, hospitals or quiet recreation (*5*, *8*, *37*). Vegetation, built form and other spatial features shape which birds persist, while the acoustic and visual traits of those birds determine what residents encounter (*1*). Sound is therefore a planning resource that can be mapped and considered alongside habitat, access and urban function (*38*). Evidence from urban parks further shows that experiences of bird sound depend on spatial setting and patterns of park use, making acoustic conditions part of landscape design (*36*). Across a city, predicted acoustic activity can identify candidates for two complementary design responses. First, seasonal acoustic zoning can identify places and periods where quiet recreation, accommodation, healthcare or teaching require particular attention. Second, ecological–acoustic co-design can arrange vegetation, open space, paths, seating and building edges to support biodiversity while offering people a wider choice of acoustic experiences. The monitoring network supplies the evidence infrastructure for both responses: it shows where these patterns are supported and where further observations are needed before intervention (*12*, *33*, *34*). Species-specific prediction locates likely intersections between biological activity and sensitive urban settings. Prediction-support classes then show where those patterns can inform screening, where local validation is needed and where monitoring must expand (*12*, *33*). Effective planning therefore requires ecological context, temporal resolution and a clear account of prediction support.

Here, we asked two connected questions. First, how does Shenzhen’s present-day urban landscape organise Asian Koel acoustic activity and its potential intersections with planning-sensitive settings? Second, where can these patterns be predicted reliably, and how can that evidence guide future urban planning and the monitoring network needed to evaluate it? We addressed these questions with an AI-assisted passive-acoustic network spanning 31 sites across Shenzhen’s urbanisation gradient in linked steps (Fig. 1). (1) We fitted a life-history-informed joint model that separated weekly acoustic opportunity, daily detectability and calling density, while carrying uncertainty from three potential-host groups into Koel estimates. (2) We mapped where predicted activity intersected planning-sensitive places and urban functions. (3) Whole-station spatial validation, the Prediction Interpretation Domain (PID) and the Area of Applicability (AOA) identified where citywide predictions were supported. (4) Coverage Expansion and Prediction Validation prioritised candidate stations in underrepresented environments and areas of high prediction uncertainty. We expected landscape and potential-host opportunity to structure the present acoustic geography, with Koel activity concentrated in seasonal and daily windows. We also expected prediction support to vary across Shenzhen, making targeted monitoring essential for extending reliable evidence. Together, these analyses connect the acoustic imprint of the present-day urban landscape with the design and evaluation of future urban spaces, recasting urban sound as evidence of how cities have been planned—and as a practical guide to how they can be planned and monitored differently.

## Results

### Potential hosts improved prediction at unmonitored sites

After accounting for the other model terms, weekly acoustic opportunity (*ψ*) declined with increasing tree cover (*β* = −1.70, 95% credible interval (CrI): −2.62 to −0.80) and built cover (*β* = −1.70, 95% CrI: −2.78 to −0.81) across the sampled urban gradient, whereas the effects of distance to water and elevation were not clearly resolved (Fig. 2A). Landscape associations differed among potential-host groups. Corvid opportunity declined with tree and built cover and increased with distance to water, while Sturnid opportunity increased with road density and declined with grass cover and elevation (Fig. 2A). Against this landscape-season baseline, potential-host opportunity added transferable ecological information. In fivefold whole-station cross-validation, the coordinate-free three-host model improved held-station expected log predictive density (ELPD) by 17.96 units relative to the matched coordinate-free model without hosts (95% site-clustered interval: 0.41 to 35.51, fig. S5B; Supplementary Data 10, Table S10C). Thus, incorporating a defining life-history background of an obligate brood parasite, namely potential-host availability, improved prediction at unmonitored sites beyond habitat and season alone. This gain concerned the three potential-host opportunities as a correlated predictive block and therefore represents their joint contribution to transfer.

**Fig. 2.**
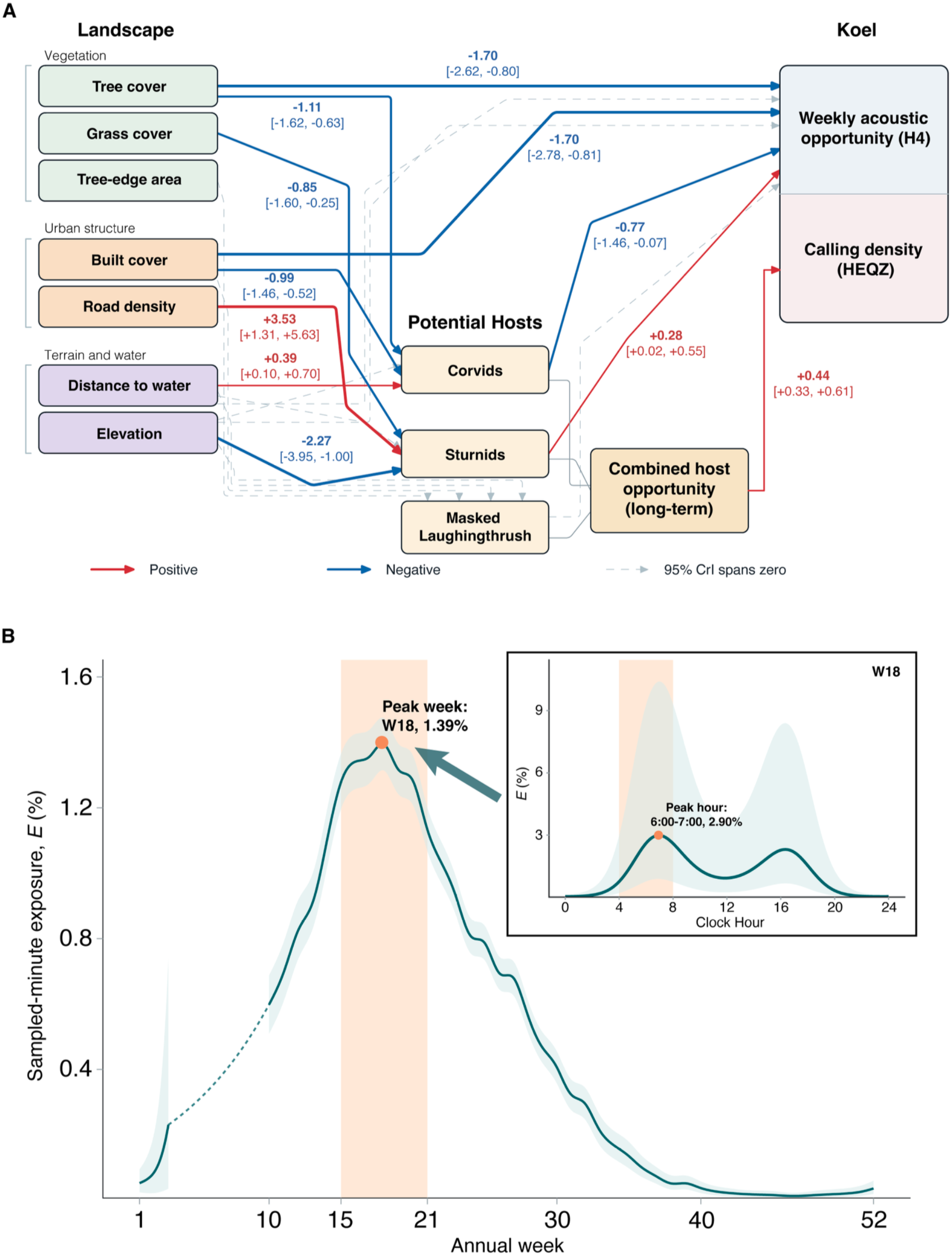
Landscape–host associations and temporal rhythms of Asian Koel acoustic activity. (A) Fitted landscape–host and host–Koel associations from 31 monitoring stations. Three separate potential-host opportunities entered the weekly acoustic-opportunity model (H4); their long-term equal-scale index entered the calling-density component (HEQZ). Red and blue arrows indicate positive and negative coefficients whose 95% credible intervals (CrIs) exclude zero; grey dashed arrows indicate intervals spanning zero. Solid grey connectors show index construction. Labels give posterior medians [95% CrIs] of standardised logit-scale coefficients. (B) Annual sampled-minute acoustic exposure, *E* = *ψpμ*^+^, at monitored sites under observed recording conditions; *ψ* is weekly acoustic opportunity, *p* is daily detectability, and *μ*^+^ is the zero-truncation-adjusted positive-day sampled-minute probability. The inset shows week 18 under the standard detection reference (144 sampled minutes per day, centred anthropogenic- and frog-event covariates, and a zero detection-level station intercept). Lines and ribbons show posterior means and 95% CrIs. The dashed section bridges unobserved weeks for visual continuity. Orange shading marks weeks 15–21 and 04:00–08:00; points mark annual and hourly peaks. Additional model summaries are provided in the Supplementary Materials.

The host-informed model left no detectable spatial pattern in held-station residuals (Moran’s *I* = 0.025, permutation *p* = 0.586), while adding broad coordinate trends did not provide a clearly resolved further gain. Matched scale comparisons likewise gave no clear evidence that either 100 m (*Δ*ELPD = −5.93, 95% interval: −13.37 to 1.52) or 500 m (−1.41, −7.24 to 4.41) improved on 250 m. We consequently retained the coordinate-free, three-host formulation at 250 m for ecological-planning prediction. Full host-model development, coordinate sensitivities, calibration and convergence diagnostics are reported in Supplementary Methods S3–S4 (Supplementary Data 8–10; figs. S3 to S5).

### Calling activity peaked in spring and at dawn

Retrieved nuisance-related posts and reports were concentrated in March and April, which accounted for 82.0% of dated social-media records and 74.1% of news records (Fig. 1B). The joint model separated three components of Koel acoustic activity: weekly acoustic opportunity (*ψ*; H4), daily detectability (*p*) and conditional calling density (*μ*; HEQZ). Daily detectability (*p*) was positively associated with recording effort, anthropogenic-event rate and frog-event rate, with all three 95% credible intervals excluding zero. These relationships confirm that daily survey effort and acoustic context shaped the probability of retaining Koel vocal evidence. Conditional calling density (*μ*) increased with the equal-scale potential-host index (posterior coefficient 0.436, 95% credible interval: 0.331 to 0.605; Fig. 2A). Adding this host term improved held-station predictive density by 92.07 ELPD units over the seasonal density model, although the gain was variable among folds (site-clustered standard error (SE) 57.17; four of five folds positive). Potential-host opportunity therefore contributed to the density of Koel-positive sampled minutes overall, but not uniformly across every spatial partition.

Adding cyclic clock-hour terms to the calling-density component (*μ*) improved held-station predictive density by 2,694.5 ELPD units (site-clustered SE 444.6) and was favoured at all 31 stations. Posterior composition translated weekly opportunity (*ψ*), daily detectability (*p*) and the positive-day sampled-minute probability (*μ*^+^) into sampled-minute acoustic exposure (*E* = *ψ* × *p* × *μ*^+^). At monitored sites under observed recording conditions, annual sampled-minute acoustic exposure peaked in week 18 (W18) at 1.394% (95% credible interval: 1.314-1.474%). Within the monitored-site annual peak week (W18), sampled-minute acoustic exposure was highest under the standard detection reference from 06:00 to 07:00, reaching 2.90% (95% CrI: 0.87–10.12%; Fig. 2B). Sampled-minute acoustic exposure therefore concentrated in spring and around dawn. Here, *E* is the probability that a scheduled sampled minute was classified as Koel-positive under the study’s recording and classification design. Full daily and hourly comparisons, posterior predictive checks and the integrated parameter-shift test are provided in Supplementary Methods S5 (Supplementary Data 11–12; fig. S6).

### Supported Koel activity expanded unevenly across Shenzhen

Citywide predictions were first separated by ecological interpretation and statistical support. Of 13,714 classified hexagonal reporting cells (HEXes), 2,712 (19.78%) formed the Primary Results and 3,033 (22.12%) were classified as Moderate Extrapolation, giving 5,745 HEXes for planning-facing summaries. A further 7,066 HEXes (51.52%) were classified as Monitoring Gaps, while 903 permanent-water HEXes (6.58%) remained Outside the Reporting Domain (Fig. 3A). Thus, 41.9% of classified HEXes were retained for planning interpretation, while 51.5% identified the principal evidence gaps in the current monitoring network. The city maps consequently show both broad seasonal expansion and a sharply uneven geography of prediction support. Citywide prediction methods, reporting populations, PID/AOA cross-tabulations and support summaries are reported in Supplementary Methods S6 (Supplementary Data 13–14; fig. S7).

**Fig. 3.**
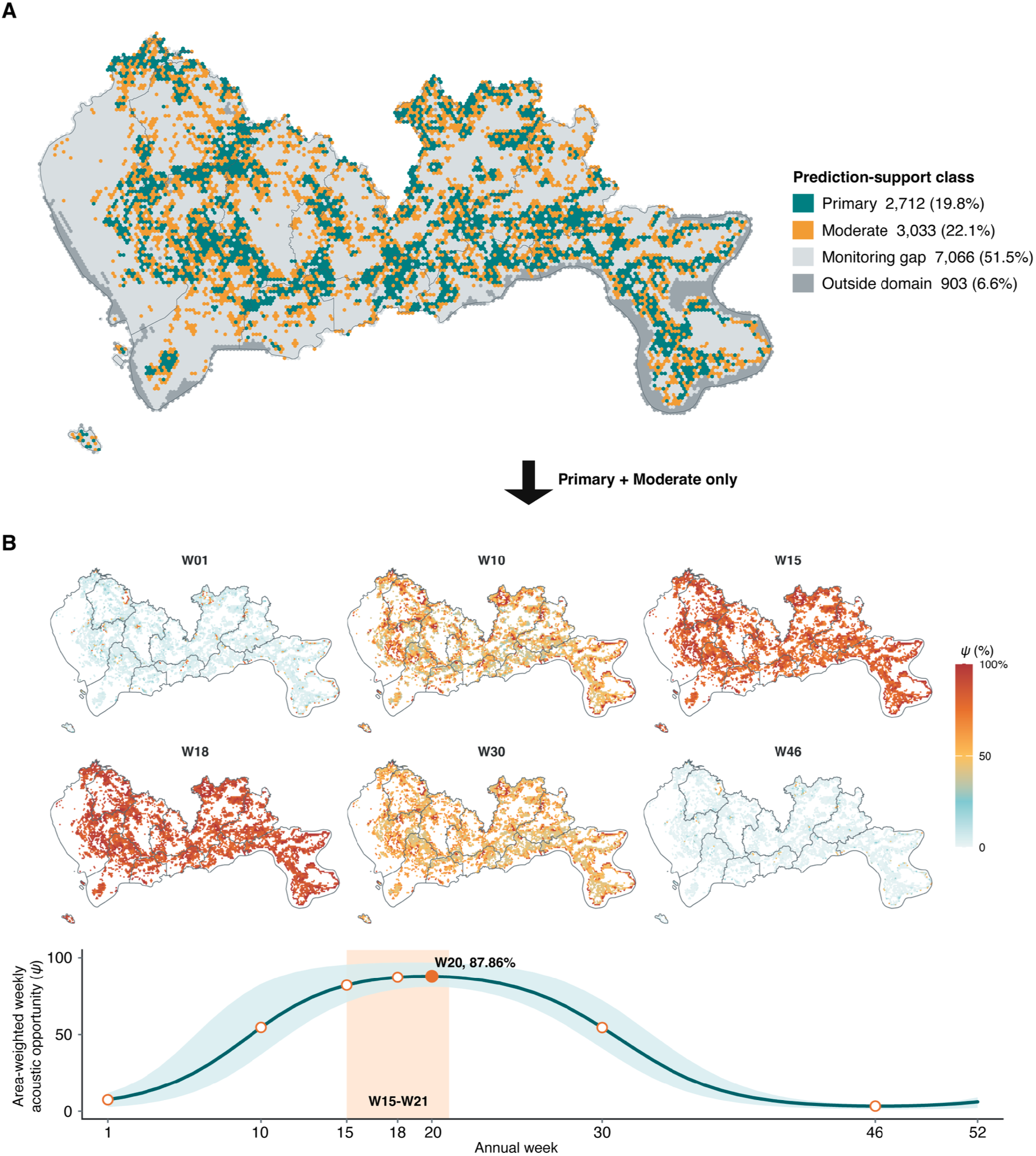
Prediction support and seasonal expansion of weekly Asian Koel acoustic opportunity across Shenzhen. (A) Cross-classification of the Prediction Interpretation Domain and Area of Applicability into Primary Results (teal), Moderate Extrapolation (orange), Monitoring Gaps (light grey) and locations Outside the Reporting Domain (dark grey), across 13,714 hexagonal (HEX) reporting cells. The corresponding legend labels are abbreviated as Primary, Moderate, Monitoring gap and Outside domain. These classes describe ecological and statistical support without rescaling prediction values; Monitoring Gaps identify environments requiring additional evidence. (B) Posterior median weekly acoustic opportunity (*ψ*) for six representative annual weeks, displayed only in Primary Results and Moderate Extrapolation HEXes. The lower curve shows the area-weighted mean across the 5,557 centroid-in-city HEXes classified as Primary Results or Moderate Extrapolation; the ribbon spans the spatial 10th–90th percentiles and describes variation among HEXes, not posterior uncertainty. The filled orange point marks the week 20 (W20) maximum (87.86%), open points mark the mapped weeks, and orange shading marks weeks 15–21 (W15–W21). Week 1 (W01) begins on 1 January. Reporting populations and support rules are provided in Supplementary Data 13–14.

Citywide projection converted this temporal concentration into a dynamic spatial pattern. Across the 5,557 centroid-in-city HEXes classified as Primary Results or Moderate Extrapolation, area-weighted weekly acoustic opportunity (*ψ*) rose rapidly through spring and peaked at 87.86% in W20 before contracting later in the year (Fig. 3B). All-day sampled-minute acoustic exposure (*E*) peaked earlier, in W17, at 1.236% (Supplementary Data 12, Table S12B). The separation between these peaks reflects the additional effects of daily detectability (*p*) and positive-day calling density (*μ*^+^): a broad weekly opportunity surface did not imply equally dense positive minutes at every place or time.

### Supported predictions overlapped planning-sensitive spaces

Within the 5,745 HEXes classified as Primary Results or Moderate Extrapolation, model-supported predictions intersected every mapped sensitive-place class and all three urban functional zones. The citywide dataset contained 3,653 points of interest (POIs) after duplicate removal, of which 763 entered these supported summaries. Across the six POI classes, mean weekly acoustic opportunity (*ψ*) during W10-W30 ranged from 68.6% to 75.5%, and peak timing fell in W20 or W21. During the W15-W21 dawn window, expected Koel-positive sampled minutes ranged from 0.226 to 0.445 per standard four-hour sampling window, while the posterior probability of at least one positive sample ranged from 15.4% to 22.0%. Together, these measures represented both seasonal acoustic opportunity and dawn activity under the reference sampling design (Table 1). Overlapping credible intervals did not support a strict ranking of receptor classes.

**Table 1.**
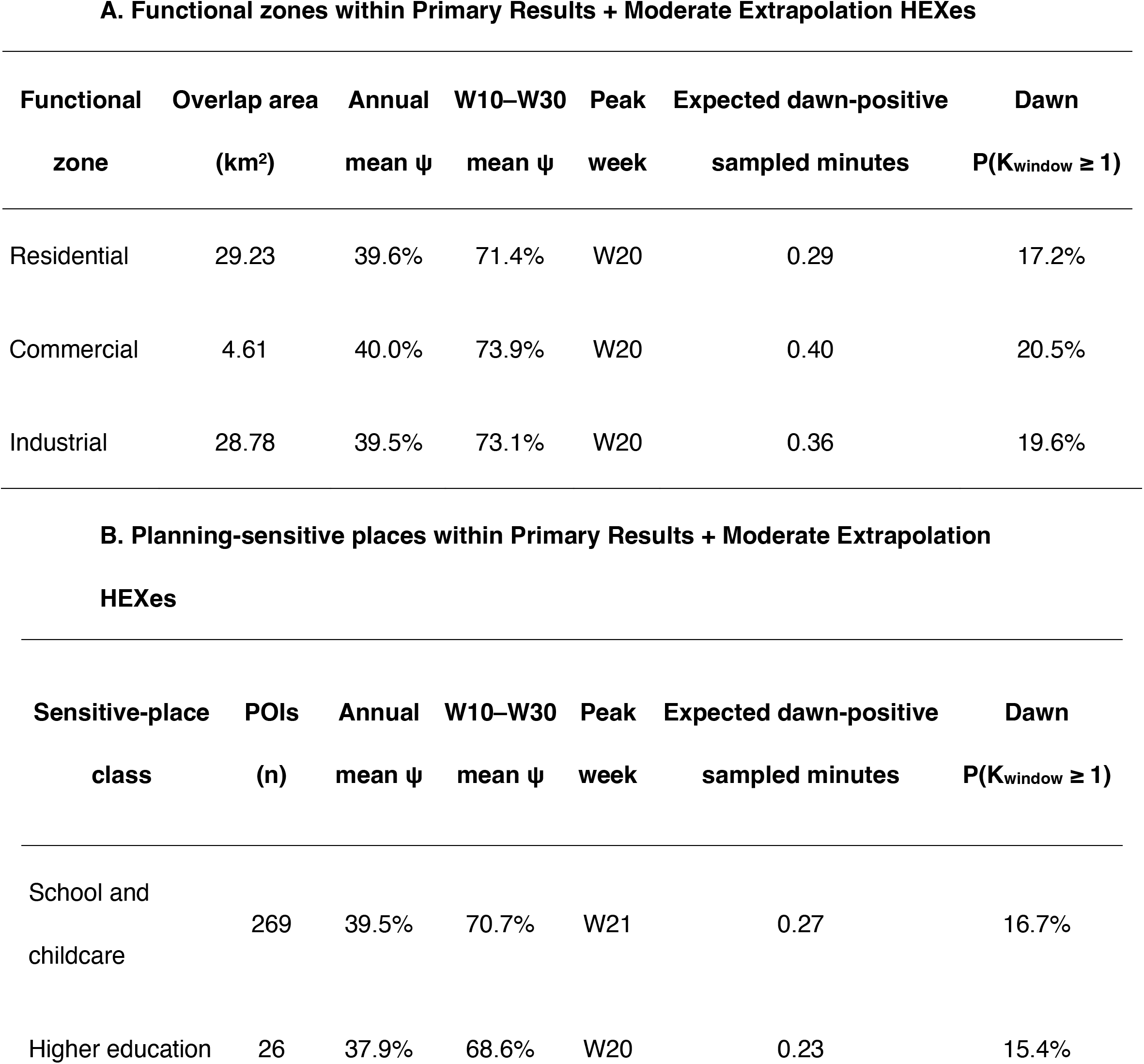
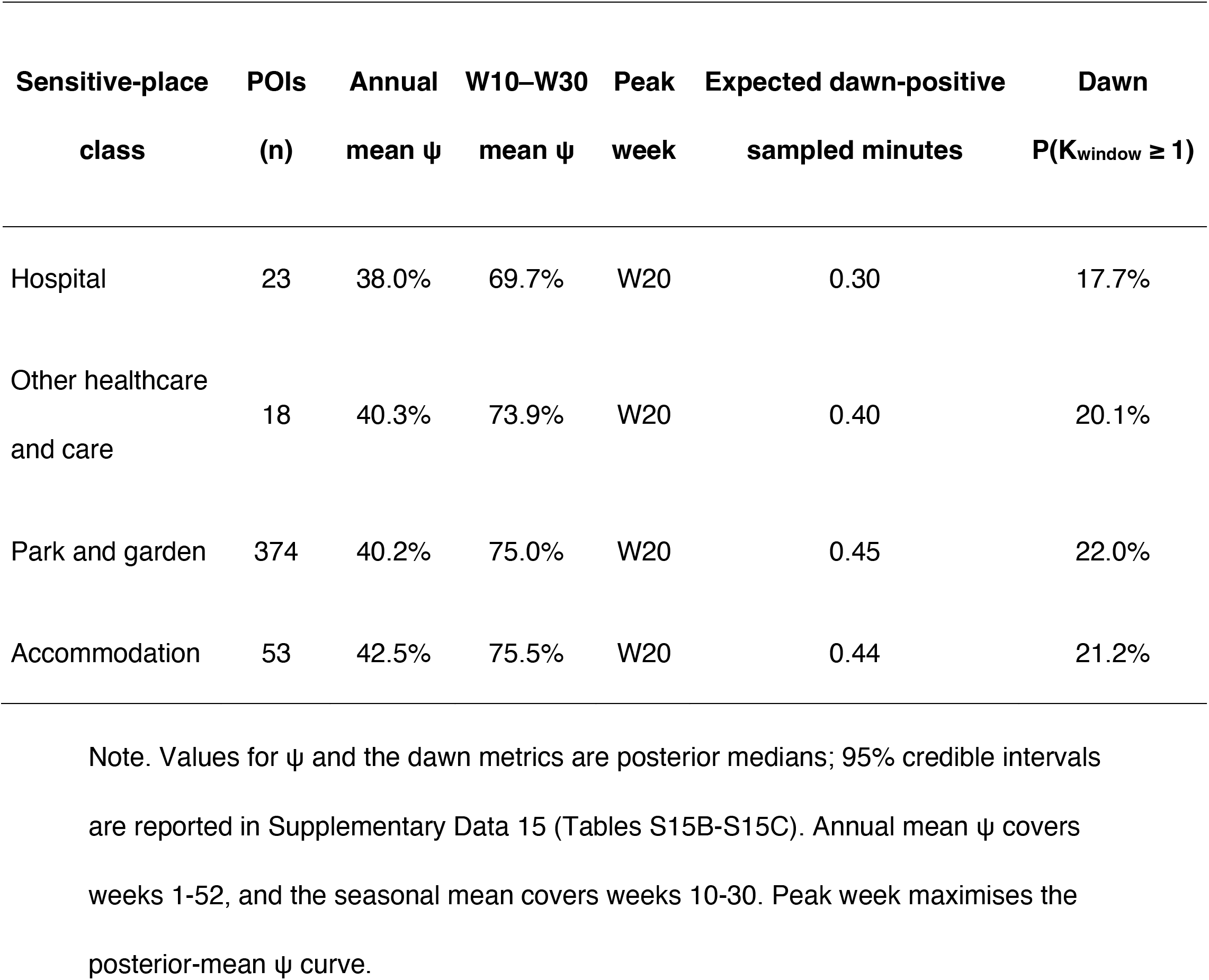
Model-supported intersections between Asian Koel acoustic activity and planning-sensitive urban spaces in Shenzhen. Summaries include hexagonal (HEX) cells classified as Primary Results or Moderate Extrapolation. Functional-zone summaries were weighted by polygon–HEX overlap area; points of interest (POIs) were equally weighted after deduplication. Annual and W10–W30 means describe group-weighted weekly acoustic opportunity (*ψ*), averaged over weeks 1–52 and 10–30, respectively. Peak week maximises the posterior-mean *ψ* curve. Dawn metrics describe the 24 scheduled one-minute samples between 04:00 and 08:00 on a reference day, averaged over weeks 15–21: the expected number of Koel-positive sampled minutes and the probability of at least one positive sample, *P*(*K*_window_ ≥ 1), where *K*_window_ is the number of positive samples. Both metrics were calculated within each posterior draw from the fitted beta-binomial process with positive-day conditioning, then averaged across weeks before group summaries were formed. Values for *ψ* and the dawn metrics are posterior medians; 95% credible intervals and complete class definitions are reported in Supplementary Data 15 (Tables S15B–S15C). These summaries represent modelled acoustic co-location under the stated sampling design.

Supported functional zone overlaps covered 29.23 km^2^ of residential, 4.61 km^2^ of commercial and 28.78 km^2^ of industrial land. Their W10-W30 mean weekly opportunities were similar (71.4%, 73.9% and 73.1%, respectively), while their corresponding dawn probabilities of at least one positive sample were 17.2%, 20.5% and 19.6% (Table 1). These results identify where seasonally concentrated Koel acoustic activity and planning-sensitive places or functions coincide. Complete class definitions, denominators and posterior intervals are reported in Supplementary Methods S7 (Supplementary Data 15; fig. S8).

Dawn metrics describe 24 scheduled one-minute samples between 04:00 and 08:00 on a reference day, averaged over weeks 15-21. K_window_ is the number of Koel-positive samples. Expected counts and P(K_window_ ≥ 1) were derived from the fitted beta-binomial process with positive-day conditioning.

Summaries include hexagonal (HEX) cells classified as Primary Results or Moderate Extrapolation. Functional zones were weighted by polygon-HEX overlap area; points of interest (POIs) were equally weighted after deduplication. These are modelled acoustic co-location summaries under the stated sampling design.

### Evidence gaps guided adaptive monitoring

The support map translated two different evidence gaps into two complementary monitoring objectives. Coverage Expansion addressed missing environmental representation by selecting Monitoring Gap HEXes that maximised prospective expansion of the Area of Applicability. Prediction Validation addressed imprecise estimates within the current interpretation domain by selecting Primary Results or Moderate Extrapolation HEXes with the widest W15–W21 weekly-opportunity (*ψ*) credible intervals. After excluding existing stations and requiring each candidate to lie at least 4.627 km from existing and earlier-selected sites—the median nearest-neighbour separation in the current network—2,734 candidate HEXes remained for Coverage Expansion and 2,017 for Prediction Validation (Fig. 4A). Mapped planning-sensitive context served only as a label or tiebreaker after these scientific scores.

**Fig. 4.**
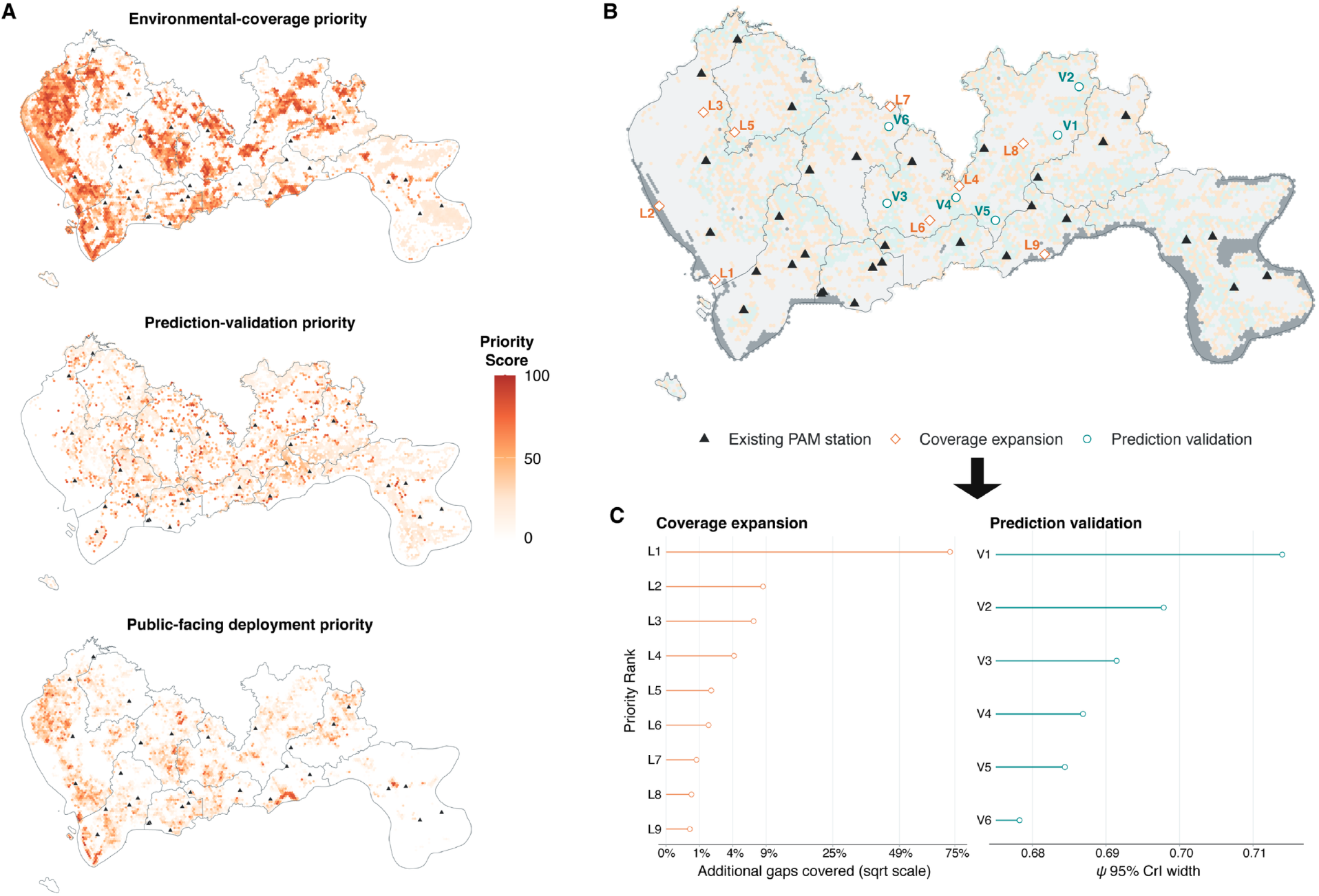
Translating prediction gaps into priorities for adaptive passive acoustic monitoring. (A) Priority surfaces for Coverage Expansion, Prediction Validation and public-facing deployment. Coverage Expansion priority measures prospective Area of Applicability (AOA) gain within Monitoring Gaps; Prediction Validation priority measures the mean W15–W21 width of the weekly-opportunity (*ψ*) 95% credible interval within Primary Results and Moderate Extrapolation areas. Public-facing deployment priority combines scientific priority with mapped planning-sensitive context as a display layer; it did not determine the selection sequences. Planning context was used only as a label or tiebreaker. Scores are scaled from 0 to 100 within each objective and quantify site-selection priority. Black triangles mark existing passive acoustic monitoring (PAM) stations. W15–W21 denotes annual weeks 15–21. (B) An illustrative 15-site portfolio comprising nine Coverage Expansion sites (orange diamonds, L1–L9) and six Prediction Validation sites (teal circles, V1–V6); labels give the sequential rank within each route. Background colours indicate Primary Results (pale teal), Moderate Extrapolation (pale orange), Monitoring Gaps (light grey) and locations Outside the Reporting Domain (dark grey). (C) Route-specific selection criteria for the selected sites. The Coverage Expansion panel shows the percentage of current Monitoring Gap hexagonal (HEX) cells newly brought within the fixed AOA threshold at each sequential addition, using a square-root x-axis; the Prediction Validation panel shows the mean W15–W21 *ψ* 95% credible-interval width on the probability scale targeted by each site. These panels report the criteria and sequential ranks used to construct the prospective portfolio. Candidate sites require field verification before deployment. Eligibility rules and complete sequences are provided in Supplementary Data 16.

The sequential design produced spatially separated candidate sites rather than a single fixed optimum. In the illustrative 15-site scenario, nine stations were allocated to Coverage Expansion to expand environmental coverage, while six were allocated to Prediction Validation to test uncertain predictions within already interpretable areas (Fig. 4B). The nine Coverage Expansion sites prospectively brought 6,884 of the 7,066 Monitoring Gap HEXes (97.4%) within the fixed Area of Applicability threshold, while the six Prediction Validation sites targeted large posterior intervals in Primary Results or Moderate Extrapolation areas (Fig. 4C). This is one transparent allocation scenario: future programmes can redistribute sites between the two objectives as deployment capacity and planning priorities change. The projected coverage gain remains prospective and does not demonstrate realised improvement in predictive accuracy. Complete candidate rankings, marginal-gain sequences and deployment scenarios are reported in Supplementary Methods S8 (Supplementary Data 16; figs. S9 to S10).

## Discussion

### Planned landscapes shape Koel acoustic activity

Shenzhen’s Koel soundscape shows how present-day urban landscape configuration organises seasonal acoustic activity across the city. Weekly Koel acoustic opportunity declined as tree cover and built cover increased, while Sturnid opportunity showed a positive conditional association within the correlated three-host block (Fig. 2A; fig. S4B). Together, these patterns point to an acoustic niche centred on the human-shaped spaces between dense woodland and intensive urban fabric: relatively open or managed green mosaics, landscaped urban edges and neighbourhoods where potential hosts remain acoustically available. Adding the three-host block improved transfer to withheld stations beyond landscape and season alone (Supplementary Data 10, Table S10C; fig. S5B). Life-history context therefore revealed a spatial dimension that generic land cover could not capture on its own: the urban acoustic pattern of a brood parasite is partly organised through the host landscape. Shenzhen’s conspicuous seasonal Koel activity can thus be read as an ecological expression of how urban habitats and bird communities have been assembled.

Urban park systems carry ecological legacies of planning (*39*). Park location, area and configuration shape contemporary bird diversity, while the surrounding landscape influences the cultural services and disservices associated with urban birds (*2*). Shenzhen extends this principle from bird communities to a species-specific seasonal signal.

Supported Koel predictions intersected parks and gardens and other planning-sensitive settings during spring, positioning urban greenspace as both biodiversity infrastructure and acoustic encounter infrastructure (Table 1; fig. S8). Landscapes created to provide recreation and contact with nature can also concentrate conspicuous wildlife signals near the places people use. In Shenzhen, the acoustic imprint of the present-day urban landscape is becoming visible: the same green network that supports urban nature also coincides with concentrated seasonal Koel activity, which may further organise where residents are most likely to encounter one of the city’s loudest seasonal birds. Our maps provide a time-point diagnosis of this pattern by showing where landscape structure, host opportunity and planning-sensitive settings converge. The next planning question is therefore clear: how can these seasonal acoustic encounters be spatially and temporally organised while retaining the biodiversity functions that produced them?

### Designing seasonal acoustic coexistence

Modelled Koel acoustic activity follows a predictable calendar, giving planners a clear seasonal window to plan for acoustic coexistence when calling is most pronounced. At monitored sites, sampled-minute acoustic exposure peaked in W18, and the W18 diel curve reached its maximum at 06:00–07:00 (Fig. 2B; Supplementary Data 12, Table S12B). Across supported city predictions, the broader W15–W21 window intersected parks and gardens, accommodation, healthcare, schools and other education settings (Table 1; fig. S8). Together, these results define a practical design window: W15–W21 and 04:00–08:00 identify periods for prioritising field checks of predicted acoustic activity and testing planning responses.

These mapped seasonal windows provide a practical basis for seasonal soundscape zoning (*38*). Within large parks and connected green networks, planners can evaluate biodiversity-listening areas, quiet-restoration areas, and alternative early-morning routes or resting spaces. The same park can then serve wildlife experience and quiet recreation through spatial and temporal organisation. Research on urban parks shows that bird sounds, traffic, landscape context and visitor characteristics jointly shape perceived soundscape quality, and that soundscape management can strengthen benefits for people and wildlife (*6*, *36*).

A second direction is ecological–acoustic co-design. Vegetation configuration, open space, paths, seating, and façades facing accommodation, healthcare or teaching facilities can be evaluated as one landscape system. In predicted spring hotspots, the maps can guide pilot projects that test where quiet-use facilities should be placed relative to the strongest predicted early-morning activity, which areas can retain seasonal birdsong as part of the intended experience, and how planting structure or circulation routes can create different acoustic experiences. Short PAM deployments before and after redesign can test whether acoustic activity shifts as expected; soundwalks and user-response surveys can reveal how those changes are experienced. This approach places acoustic experience alongside habitat, cooling, recreation and visual quality as a normal dimension of multifunctional greenspace planning (*10*). Spatial configuration, activity scheduling and seasonal communication therefore provide immediate planning tools, followed by targeted habitat adjustments when local evidence identifies a clear need. Prediction support determines where these actions begin: supported areas can enter design screening, and weakly represented environments first receive additional monitoring.

### Prediction support as planning infrastructure

Prediction support turns the citywide map into planning infrastructure. Supported maps provide a screening layer for identifying where and when seasonal acoustic activity may intersect planning-sensitive urban settings, and where field validation is required before intervention. The Prediction Interpretation Domain (PID) identifies land-cover contexts for ecological interpretation, while the Area of Applicability (AOA) tests whether a HEX resembles environments represented by the 31-station network. Their cross-classification assigns a planning action to every location. Primary Results can enter seasonal screening and design; Moderate Extrapolation identifies areas for field validation; Monitoring Gaps mark environments where new observations should come first; and locations Outside the Reporting Domain remain excluded (Fig. 3A; Supplementary Data 13, Table S13B; Supplementary Data 14, Table S14B; fig. S7C). This hierarchy shows where the spring and dawn zones proposed above can guide design, where they provide hypotheses to test, and where the evidence base must expand. Because 51.5% of Shenzhen’s mapped grid fell into Monitoring Gaps, missing evidence is itself a major planning result (Fig. 3A; Supplementary Data 14, Table S14A) (*12*).

The same framework specifies how the city should learn next. Coverage Expansion selects sites that extend environmental representation into Monitoring Gaps (Fig. 4A; fig. S9A). Prediction Validation targets Primary Results and Moderate Extrapolation areas where W15–W21 weekly acoustic opportunity remains most uncertain (Fig. 4A; fig. S9B). The first route enlarges the city that can be interpreted; the other tests uncertain spring predictions inside the current domain. In the illustrative 15-site portfolio, nine Coverage Expansion sites would prospectively bring 6,884 of 7,066 Monitoring Gap HEXes (97.4%) within the fixed AOA threshold, while six Prediction Validation sites would test locations with wide spring credible intervals (Fig. 4B–C; Supplementary Data 16, Tables S16B– S16C; fig. S10 (A to C)). Planning-sensitive places and urban functions supply deployment context once scientific priority has been established (fig. S9C).

This design converts citywide prediction from a static endpoint into an adaptive decision interface. Every HEX points to an action—screen, validate, monitor or exclude—and every new PAM deployment is chosen for the evidence it can add. Repeated updating can expand reliable coverage, challenge uncertain predictions and keep soundscape planning aligned with a changing city (*33*).

### Future outlook: from acoustic maps to adaptive landscapes

Once prediction support is linked to monitoring design, the city can begin to learn from each planning cycle. Our maps provide a time-stamped diagnosis of how Shenzhen’s existing landscape is associated with seasonal Koel acoustic activity and its potential intersections with planning-sensitive settings. The next monitoring rounds can turn park renewal, planting reconfiguration, route separation and seasonal soundscape zoning into testable interventions. PAM deployed before and after design changes can update weekly opportunity *ψ*, detectability *p* and calling density *μ*, while soundwalks, visitor surveys and complaint records can measure how the same acoustic conditions are experienced. These linked observations would reveal which configurations sustain wildlife activity, which provide valued listening opportunities and which concentrate unwanted early-morning encounters. The resulting sequence—planning decision, wildlife response, public experience, re-monitoring and redesign—makes landscape planning a cumulative learning process (*34*).

The same structure can travel beyond the Koel and beyond Shenzhen. For brood parasites, potential-host opportunity supplies the life-history layer; for other planning questions, this layer may describe nesting resources, food, roosts, wetlands or movement corridors. The transferable architecture remains consistent: AI-assisted observations, explicit correction for detectability, behaviour-specific activity estimates, validation at unseen sites, mapped prediction support and evidence-guided redeployment. Repeating this cycle across projects would create an urban acoustic observatory that records how planned landscapes reorganise encounters between wildlife and people through time. The next evidence layer should connect modelled activity with sound level, human presence and perceived disturbance, allowing cities to evaluate ecological function and public experience together. Acoustic maps can then operate as renewable planning evidence, updated as landscapes change and used to guide the next intervention, the next survey and the next design (Fig. 1C).

Present-day urban landscape configuration helps organise when and where wildlife becomes audible in everyday life. In Shenzhen, AI-assisted acoustic monitoring showed that Koel activity emerged from the joint geography of landscape configuration, potential hosts and season, then concentrated into spring and early-morning encounters across planning-sensitive spaces. Distinguishing planning-ready predictions from areas requiring new observations turned the resulting maps into actionable evidence. Cities can use this evidence to diagnose the audible ecological consequences of earlier urban design, establish seasonal soundscape zones, coordinate biodiversity and quiet-use functions, and target future monitoring. By integrating life-history context, spatial validation and adaptive sampling, our framework offers a practical route for designing urban landscapes where wildlife activity and human acoustic experience can be understood, anticipated and managed together.

## Materials and Methods

### Experimental design and acoustic monitoring

Shenzhen is a subtropical megacity in southern China spanning densely built districts, urban greenspaces and remnant evergreen forest. Thirty-one passive acoustic monitoring stations sampled this gradient (Fig. 1A; Supplementary Methods S1.1; Supplementary Data 1; fig. S1). The analysis focused on Asian Koel (*Eudynamys scolopaceus*) acoustic activity. Each station used an AudioMoth recorder mounted approximately 1.5 m above ground, sampling nominally for one minute every 10 min at 48 kHz and 16-bit resolution (*40*).

Monitoring was conducted from 31 March 2024 to 12 August 2025. We retained 9,912 effort-confirmed station-day surveys, including 2,682 Koel-positive days, representing 1,398,905 recorded sampled-minute opportunities and 1,500 station-weeks within 68 indexed sampling weeks (Supplementary Methods S1.2; Supplementary Data 2; fig. S1). Corresponding dates from both years informed a common annual cycle, with unobserved intervals retained as gaps. Recordings were downsampled to 32 kHz and processed sequentially by sound-event detection, spectrogram localisation and a locally adapted BirdNET v2.4 classifier. Records were retained when event-detection and species-identification confidence were both ≥ 0.5 (Supplementary Methods S1.3; Supplementary Data 3). Days with verified effort but no retained Koel record were coded as non-detections; days without usable effort remained missing. These histories quantify Koel vocal evidence. The complete analytical workflow is summarised in Fig. 1C. To describe the public context of Koel-related disturbance, we also summarised social-media posts and supplementary news reports published from January 2023 to September 2026 by calendar month (Fig. 1B; Supplementary Methods S1.4).

### Weekly acoustic opportunity, potential hosts and spatial validation

Daily non-detections could indicate either no weekly Koel vocal activity or failure to record it. For station *i*, week *t* and day *j*, we modelled

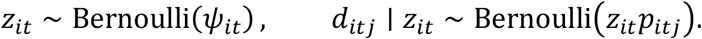

Here, *d_it_*_j_ = 1 when surveyed day *j* at station *i* in week *t* contained at least one retained Koel-positive sampled minute, and 0 otherwise; *z_it_* is an unobserved binary indicator of whether station *i* contained Koel acoustic activity during week *t*; *ψ_it_* is the probability of that weekly opportunity; and *p_it_*_j_ is the probability of retaining at least one Koel-positive minute on surveyed day *j* when *z_it_* = 1 (*31*, *32*). Detectability varied with recording effort, anthropogenic-event rate and frog-event rate, with a detection-level station intercept for persistent differences in recording and classification conditions.

These covariates described daily survey effort and the acoustic observation environment (Supplementary Methods S2; Supplementary Data 4–5; fig. S2).

The retained 250-m Koel model included tree cover, built cover, distance to recurrent water, elevation and annual sine and cosine terms. Dynamic World V1, JRC Global Surface Water v1.4 and NASA SRTMGL1 v3 supplied these variables (*41–43*); a dated OpenStreetMap snapshot supplied road variables for potential-host models (Supplementary Methods S3; Supplementary Data 6–7). The three detection-corrected host responses were a four-species Corvid assemblage, a three-species Sturnid assemblage and Masked Laughingthrush (*Pterorhinus perspicillatus*) (Supplementary Methods S3; Supplementary Data 8; fig. S3). Each potential-host response was estimated with its own detection-corrected landscape-season model. Posterior draws from the three host models then entered a joint transfer model without projected-coordinate covariates (H4-P0 in Supplementary Data 9, Table S9C; hereafter, the primary three-host transfer model), summarised schematically as:

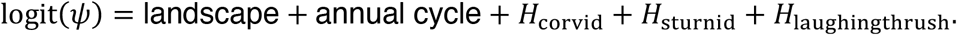

The full coefficient-level formula is provided in Supplementary Methods S4.5. Potential-host acoustic opportunity quantified the predicted weekly vocal activity of each potential-host group. The three opportunities entered the transfer model as one correlated predictive block, and their coefficients collectively represented the contribution of host context to Koel prediction. We propagated host uncertainty through three independently sampled host-opportunity surfaces per validation fold and pooled the resulting Koel posterior predictions (Supplementary Methods S4.4). The primary transfer model omitted the occurrence-level station intercept because site-specific values are unavailable at unmonitored locations. Its detection-level station intercept was retained; predictions for withheld or new sites averaged over the fitted among-station distribution of this observation effect (Supplementary Methods S4.1–S4.5; Supplementary Data 9; fig. S4).

Model development followed five prespecified stages: selection of the daily detection structure, fitting of the three potential-host models, selection of the Koel landscape– season base model, comparison of predefined host structures, and whole-station spatial validation. The complete model formulae and selection sequence are provided in Supplementary Methods S2–S4 and Supplementary Data 4–10. Fivefold cross-validation withheld complete stations; scaling and fitting used training stations only, and complete daily histories were scored at test stations. Held-station expected log predictive density (ELPD) was the primary criterion for comparing transfer to withheld stations, with Brier score, calibration, Markov chain Monte Carlo (MCMC) diagnostics and residual Moran’s *I* as supporting evidence (Supplementary Methods S4.4–S4.6; Supplementary Data 10). Because landscape relationships can depend on the spatial extent of measurement, matched scale sensitivity used the same no-host coordinate-trend structure at 100, 250 and 500 m (*44*); neither alternative radius clearly improved held-station transfer. We therefore retained a 250-m neighbourhood for subsequent modelling and citywide prediction. At that scale, projected-coordinate terms did not yield a clearly supported gain for the primary three-host transfer model, and the coordinate-free model showed no detectable residual spatial autocorrelation (Supplementary Methods S4.4–S4.7; Supplementary Data 10; fig. S5).

### Joint calling density and sampled-minute acoustic activity

Daily detection and non-detection sequences identified weekly acoustic opportunity. To quantify calling density on detected days, we additionally modelled *K_it_*_j_, the number of Koel-positive sampled minutes among *N_it_*_j_ recorded minutes:

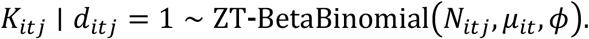

Weekly acoustic opportunity, daily detectability and the beta-binomial count parameter *μ* were estimated in one daily likelihood (*45*). All surveyed days informed *ψ* and *p*, whereas positive-day counts additionally informed *μ*; missing days contributed neither detections nor counts. The host-informed calling-density component (HEQZ) related *μ* to the annual cycle and a site-level equal-weight index of annual-mean potential-host opportunity, constructed after separate standardisation of the three host responses; the same host-posterior draw sets propagated uncertainty (Supplementary Methods S5.1– S5.2; Supplementary Data 11).

To resolve diel variation, we retained every recorded hour within positive days, including zero-count hours. We selected the hourly calling-density structure by withholding complete stations and comparing three prespecified candidates: an annual-only model, a model adding two cyclic clock-hour harmonics, and a model adding season-by-hour interactions. The retained harmonic model supplied hourly structure through modular posterior composition. A confirmatory integrated fit showed that the added hourly information produced no material shift in the shared *ψ* or *p* parameters (Supplementary Methods S5.3–S5.4; Supplementary Data 11; fig. S6).

Posterior draws were combined before calculating sampled-minute acoustic exposure *E*, expected Koel-positive sampled minutes and the probability of at least one positive sampled minute in a time window. The count model was fitted to positive days, so its beta-binomial mean *μ* describes the underlying count process before conditioning on a positive daily total. We converted *μ* to the zero-truncation-adjusted sampled-minute probability *μ*^+^, which conditions the minute-level probability on a model-positive day, and calculated *E* = *ψ* × *p* × *μ*^+^. Thus, *E* is the posterior probability that a scheduled sampled minute would be classified as Koel-positive under the stated observation conditions. Window-level detection probabilities were calculated for each posterior draw from beta-binomial zero-count probabilities, adjusted for positive-day conditioning and combined with weekly acoustic opportunity and daily detectability (Supplementary Methods S5.5; Supplementary Data 12; fig. S6).

### Citywide prediction and interpretation support

The selected primary three-host transfer model was projected to 13,714 fixed hexagonal reporting cells (hereafter, HEX cells) across 52 annual weeks. Environmental predictors were calculated within 250-m-radius circles around HEX centroids, matching the fitted ecological neighbourhood; HEX polygons stored and displayed predictions but did not define that neighbourhood. Potential-host and Koel posterior draws were propagated before weekly opportunity and sampled-minute acoustic activity were summarised (Supplementary Methods S6.1; Supplementary Data 13).

We attached two independent support labels that answered different questions. The Prediction Interpretation Domain (PID) asked whether a HEX provided a clear terrestrial context for ecological interpretation and classified its Dynamic World land-cover context as PASS, UNCERTAIN or EXCLUDED, with categorical permanent water identified from ESA WorldCover 2021 v200 (*46*). The Area of Applicability (AOA) asked how closely its seven-variable environmental setting resembled those represented by the 31 monitoring stations and classified statistical support as SUPPORTED, BOUNDARY, EXTRAPOLATED or SEVERELY UNSUPPORTED using training-range checks and nearest-station dissimilarity. Detailed inputs, deterministic rules and thresholds are provided in Supplementary Methods S6.2–S6.3 and Supplementary Data 13–14. We cross-classified PID and AOA as Primary Results, Moderate Extrapolation, Monitoring Gaps or Outside the Reporting Domain. Primary Results and Moderate Extrapolation formed the planning-reporting population; Monitoring Gaps identified reportable environments that were poorly represented by the network; categorical permanent water remained Outside the Reporting Domain. Both labels accompanied predictions without changing or rescaling their values (Supplementary Methods S6.4; fig. S7).

### Overlap with sensitive places and urban functions

Planning summaries were restricted to Primary Results and Moderate Extrapolation HEXes. A dated OpenStreetMap snapshot supplied six classes of sensitive places and residential, commercial and industrial polygons. Deduplicated points of interest (POIs) were assigned to their containing HEX and weighted equally, whereas functional zones were weighted by polygon-HEX overlap area (Supplementary Methods S7.1–S7.3; Supplementary Data 15).

For reporting, annual prediction weeks were denoted W01–W52 on a 52-week prototype beginning on 1 January. Observation dates were assigned to 7-day day-of-year bins, with the final bin capped at W52; this reporting axis was separate from the dated seasonal basis used for fitting. For each urban group, annual mean weekly acoustic opportunity (*ψ*) represented year-round Koel vocal activity, while the W10–W30 mean and peak week summarised the magnitude and timing of the broad breeding-season acoustic pulse. Matched *ψ*-, *p*-and *μ*^+^-posterior draws were then summarised at the window scale. Expected Koel-positive sampled minutes and the probability of at least one positive sample, *P*(*K*_window_ ≥ 1), were calculated for the 24 scheduled samples in a reference 04:00–08:00 window and averaged across W15–W21. Together, these metrics located where seasonal Koel acoustic activity intersected planning-sensitive places and urban functions (Supplementary Methods S7.4; Supplementary Data 15; fig. S8).

### Adaptive monitoring design

Candidate HEXes had PASS or UNCERTAIN PID status, complete values for the seven environmental predictors fitted across the Koel and potential-host models, and no existing station. New sites were kept at least 4.627 km from every existing and previously selected station. This distance equals the median nearest-neighbour separation among the 31 current stations and limits redundant spatial clustering (Supplementary Methods S8.1; Supplementary Data 16). We used two complementary routes. Coverage Expansion sequentially selected Monitoring Gap HEXes that maximised prospective coverage of the monitored environmental space. Prediction Validation selected Primary Results and Moderate Extrapolation HEXes with the greatest mean width of the weekly-opportunity 95% credible interval across W15–W21 (Supplementary Methods S8.2–S8.3; Supplementary Data 16; fig. S9). Scientific scores determined the sequence; mapped sensitive places and urban functions described deployment context and resolved exact ties. We assembled an illustrative 15-site portfolio and retained field checks for access, permission, security and local recording conditions (Supplementary Methods S8.4; Supplementary Data 16; fig. S10).

### Computation and reproducibility

Environmental processing used Google Earth Engine and projected spatial workflows in R (*47*). Occupancy models were fitted with spOccupancy; daily and hourly hurdle models were fitted in Stan through cmdstanr (*48*, *49*). OpenStreetMap data were processed with osmium-tool and R spatial operations (Supplementary Methods S9.1; Supplementary Data 17, Table S17A). Software versions, priors, random seeds, sampling settings and reproducibility checks are reported in Supplementary Methods S9.2–S9.3 and Supplementary Data 17 (Tables S17B–S17C).

### Statistical analysis

All statistical analyses were conducted in R 4.5.2. Model estimates and derived acoustic metrics were summarised using posterior distributions and 95% credible intervals. Convergence was assessed using *R*^^^ and bulk and tail effective sample sizes. Stan fits were additionally checked for divergent transitions and maximum-treedepth hits. Paired differences in held-station predictive performance were aggregated at the station level, with uncertainty reported using station-clustered intervals. Model-fitting settings and diagnostic thresholds are reported in Supplementary Methods S9.2 and Supplementary Data 17 (Table S17B); validation procedures are described in Supplementary Methods S4.4–S4.7.

## Supporting information

Supplementary Data 1. Monitoring stations and survey coverage

Supplementary Data 2. Recording effort and detection-history construction

Supplementary Data 3. Acoustic-identification workflow

Supplementary Data 4. Daily-detection model comparison

Supplementary Data 5. Daily-detection model checks

Supplementary Data 6. Model variables and reporting quantities

Supplementary Data 7. Spatial data sources and processing

Supplementary Data 8. Potential-host model development and validation

Supplementary Data 9. Koel acoustic-opportunity model development

Supplementary Data 10. Spatial-transfer validation and scale sensitivity

Supplementary Data 11. Joint acoustic-model components and calling density

Supplementary Data 12. Acoustic-metric definitions and reference summaries

Supplementary Data 13. Citywide projection and prediction-support rules

Supplementary Data 14. Citywide prediction-support summaries

Supplementary Data 15. Acoustic activity in planning-sensitive urban settings

Supplementary Data 16. Additional monitoring-site selection

Supplementary Data 17. Software, model settings and reproducibility records

Description of Additional Supplementary Files

## Acknowledgements

We thank the members of the Avian Ecology and Evolution Group at Sun Yat-sen University and the Sounds Great acoustic discussion group for their support and helpful discussions throughout this study. We are especially grateful to Xuecong Zhang, Guancheng Zhu and Guangpeng Wei for their assistance with deploying the acoustic recorders.

## Author contributions

Y. C.: Conceptualization, Formal analysis, Investigation, Writing – original draft.

Z. H.: Conceptualization, Investigation, Writing – review & editing.

C. Z.: Methodology, Software, Formal analysis, Investigation.

Z. L.: Conceptualization, Formal analysis, Writing – review & editing.

L. Z.: Conceptualization.

Z. C.: Writing – review & editing.

W. F.: Writing – review & editing.

W. L.: Conceptualization.

T. Z.: Supervision, Funding acquisition, Writing – review & editing.

Y. L.: Conceptualization, Supervision, Funding acquisition, Writing – review & editing.

## Data, code, and materials availability

Numerical results are provided in the main text and Supplementary Materials, including Supplementary Data 1–17. Analysis code, de-identified model inputs and selected posterior draws and summaries are available at https://github.com/Fly-Carrot/urban-koel-acoustic-planning. The repository documentation describes the released files, data sources and access conditions.

