## Supplementary Information (Methods and Figs. S1-S10) for "Urban planning leaves an audible legacy in a megacity’s bird soundscape"

### Supplementary Methods S1. Passive acoustic monitoring and media records

#### S1.1 Monitoring sites and recording schedule

The monitoring network comprised 31 stations distributed across Shenzhen, southern

China (22.504–22.827° N, 113.840–114.566° E). Stations spanned densely built districts, urban greenspaces and remnant evergreen forest, thereby sampling the principal landscape contexts represented in the study. Station coordinates, survey coverage, response summaries and spatial-validation fold membership are reported in Supplementary Data 1. The spatial distribution of monitoring stations, fixed validation folds and daily recording effort are shown in fig. S1.

AudioMoth recorders were mounted on trees approximately 1.5 m above ground (*40*). Locations were chosen to represent the surrounding landscape while avoiding immediately adjacent loudspeakers, pump houses and other persistent mechanical noise sources that could saturate recordings. Recorders sampled nominally one minute every 10 min at 48 kHz and 16-bit resolution, giving up to 144 scheduled sampled-minute opportunities per complete station-day. Coordinates were recorded in WGS 84 and transformed to China Geodetic Coordinate System 2000 / 3-degree Gauss–Krüger CM 114° E (EPSG:4547) for distance-based spatial analysis. The 31 stations formed a fixed environmental-gradient network designed to span major landscape contexts. Because site selection prioritised environmental coverage, citywide inference was conditioned on the environmental and geographic support represented by the monitored network.

**S1.2 Which recording days entered the analysis**

Monitoring covered 31 March 2024–12 August 2025. Survey effort was reconstructed from the audio archive independently of whether any bird was detected. Each station-date combination was classified as an effort-confirmed survey or as missing. A day was retained whenever usable recordings established genuine sampling effort, including lower-effort days and days on which no bird event was retained. A day without verifiable usable effort remained missing and was never converted to a non-detection.

The station-by-week response array contained 14,756 potential daily survey slots (31 stations × 68 indexed weeks × 7 days). Of these, 9,912 were effort-confirmed and 4,844 were missing. The valid surveys comprised 2,682 Koel-positive days and 7,230 non-detection days, including 350 effort-confirmed days with no retained bird event of any species. They represented 1,398,905 minute-indexed recording opportunities entering the fitted response and 1,500 station-weeks within 68 indexed sampling weeks. These quantities, their arithmetic reconciliation and the distinction between the scheduled sampling design and usable audio are given in Supplementary Data 2.

For the weekly occurrence models, annual seasonality was calculated from the actual start date of each observed week on a continuous 365.2425-day annual clock. Observations from different years were thereby aligned to the same annual cycle while retaining their actual dates and intervening sampling gaps. The 68-week identifiers preserved their actual calendar dates and intervening gaps; they were not treated as an uninterrupted 68-week sequence. To estimate a shared annual cycle across years, each observed week was mapped to an annual clock using its calendar day of year:

$$\sin_t = \sin\left(2\pi \frac{\mathrm{DOY}_t - 1}{365.2425}\right), \qquad \cos_t = \cos\left(2\pi \frac{\mathrm{DOY}_t - 1}{365.2425}\right).$$

Thus, observations from corresponding dates in 2024 and 2025 informed the same seasonal curve, whereas unobserved intervals contributed no artificial survey occasions. In particular, the 42-day gap between week starts on 20 January and 3 March 2025 remained an explicit absence of sampling.

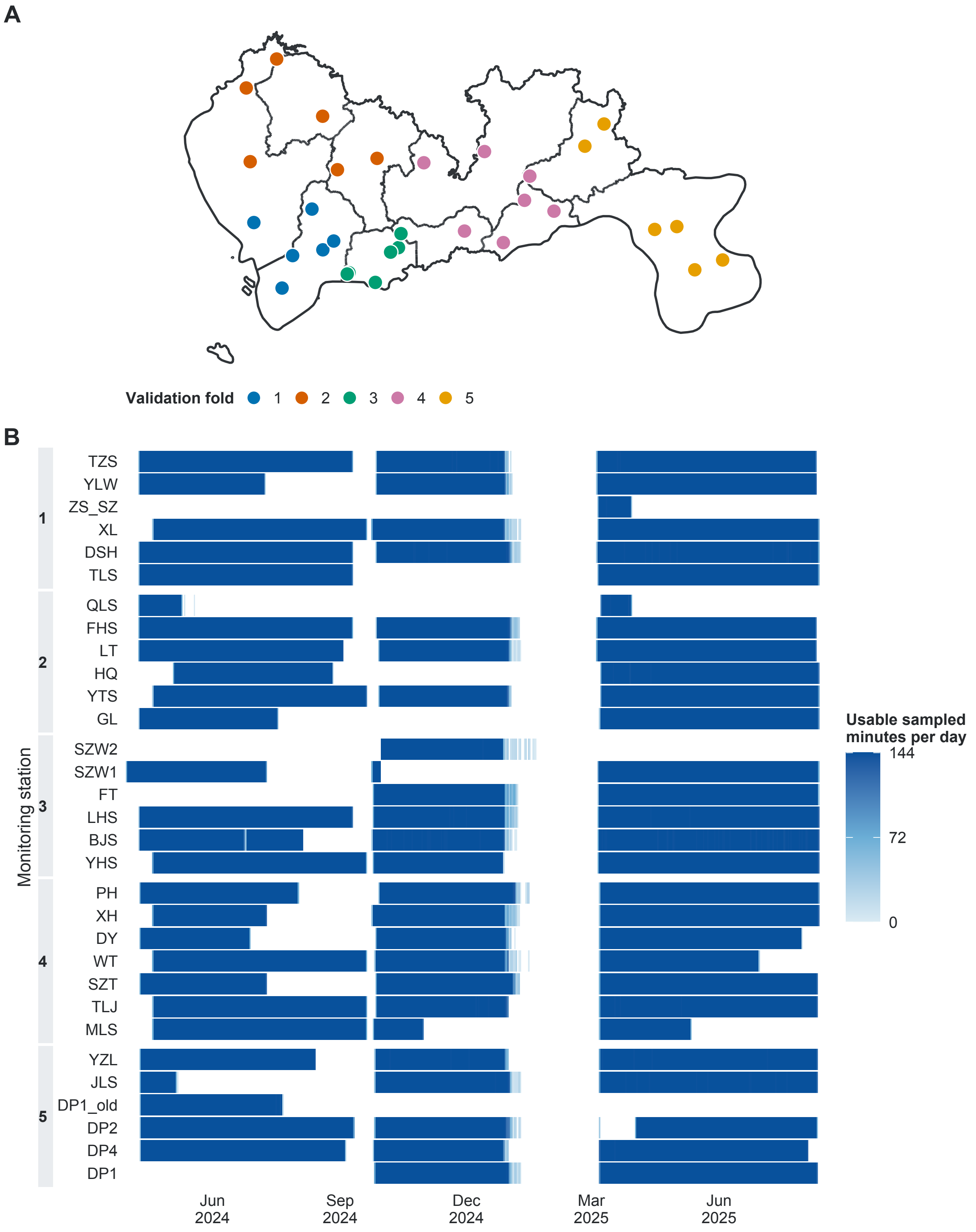


Fig. S1. Spatial distribution of passive acoustic monitoring stations and temporal recording effort. (A) Locations of the 31 monitoring stations in Shenzhen. Colours indicate the fixed whole-station spatial-validation folds used throughout model and scale comparisons; district boundaries are shown for reference. (B) Daily recording effort from 31 March 2024 to 12 August 2025, expressed as the number of sampled-minute opportunities retained for analysis, with a nominal maximum of 144 per complete station-day. White cells indicate days without usable recording effort, rather than Koel non-detections, and observed temporal gaps were retained without interpolation. Station rows are grouped by spatial-validation fold. Coordinates are reported in WGS 84; metric spatial analyses used EPSG:4547.

#### S1.3 How Koel calls were identified and counted

Each one-minute recording was downsampled to 32 kHz and divided into six 10-s segments. Three automated models then performed different tasks in sequence.

1. Finding candidate sound events. A detector combining self-supervised ATST-Frame representations with an EACRNN classifier was trained through a two-stage, semi-supervised Mean Teacher strategy using strongly labelled, weakly labelled and unlabelled audio (*50*, *51*). It returned frame-level boundaries for urban sound classes, including birds, insects, frogs and anthropogenic sounds. Evaluation across confidence operating points yielded polyphonic sound detection scores of PSDS1 = 67.64% and PSDS2 = 91.05%; PSDS1 imposed stricter temporal-overlap requirements than PSDS2. On a separate, visually checked Guangdong bird-event set, macro- and micro-recall were 75.85% and

75.91%.

2. Locating bird sounds within spectrograms. Bird-positive segments were converted to short-time Fourier-transform spectrograms using a 2,048-sample window and 512-sample hop. A YOLOv8n-based (https://github.com/ultralytics/ultralytics) detector trained from manually drawn time-frequency boxes mapped each signal to onset, offset and frequency limits. The development data contained 4,162 spectrograms from 177 species (3,329 training; 833 validation), all labelled as one "bird" class because this stage localized signals rather than identified species. The evaluated YOLOv8n+PA+DSConv variant achieved mAP50 = 67.3%, scored at a box intersection-over-union threshold of 0.5.

3. Assigning species names. A BirdNET v2.4-derived classifier was adapted to Guangdong birds through transfer learning and semi-supervised refinement with local unlabelled recordings. Its training corpus contained 67,970 strongly labelled 3-s clips from 579 species, assembled from Xeno-canto, BirdCLEF and local recordings after automated screening, deep-embedding clustering and manual review. On a balanced, independently labelled 50-species local test set, top-1 accuracy was 75.75% without confidence filtering and 89.0% among predictions with species confidence at least 0.5.

For this analysis, a candidate Koel record was retained only when both sound-event-detection and species-identification confidence were at least 0.5. Names were reconciled against the regional checklist and a curated taxonomic dictionary. Supplementary Data 3 summarizes model roles, training evidence, validation scope, thresholds and outputs. These performance values come from shared multi-species evaluations; no independent Koel-specific sensitivity or specificity estimate was available.

Retained outputs were aggregated by station and date. For every effort-confirmed survey we recorded the number of usable sampled-minute opportunities, the number containing at least one retained Asian Koel event and whether the day contained any Koel-positive minute. The final dataset contained 10,127 Koel-positive sampled minutes across 2,682 positive station-days. Effort-confirmed days with zero positive minutes were retained as non-detections; days without verified effort remained missing.

Up to seven daily records formed each station-week detection history. These histories supplied three linked pieces of evidence to the hierarchical model: whether weekly acoustic opportunity was present, whether it was detected on a surveyed day and, on positive days, how densely Koel-positive minutes occurred. Their probabilistic definitions are given in Supplementary Methods S2 and S5. All three outcomes quantify recorded vocal activity. They do not identify abundance, continuous residence, territory ownership, nesting or general habitat use.

#### S1.4 Social-media and news records

We compiled social-media posts returned by searches for “Shenzhen AND Asian Koel” in Chinese on Weibo and RedNote, together with supplementary online news reports about Koel-related disturbance in Shenzhen. The publication-date window was 1 January 2023–30 September 2026. Screening used saved search text for Weibo, titles or available descriptions for RedNote, and saved article text for news. Each record was assessed separately through manual checking and classified as a personal complaint or self-reported disturbance (A), relevant reporting, retelling or discussion (B), other Koel content (C), unrelated content (D), or insufficient information (E). Shenzhen relevance was recorded separately; city-specific search results without an explicit location remained flagged as uncertain.

The monthly summaries retained categories A and B with known publication dates: 712 social-media records (Weibo, 675; RedNote, 37) and 27 news records. Four social-media A/B records without known dates were excluded. Each source-specific record identifier was counted once; reposts and separate news publications could contribute separate records. We first summed raw A and B counts across the two social-media platforms by calendar month, then divided each monthly category count by the group's total dated A+B records. News percentages were calculated separately in the same way. Each group therefore sums to 100% across both categories and all 12 months. January–September pooled four years, whereas October–December pooled three because the window ended in September 2026. These summaries describe publication patterns in the retrieved material and provide public context for the acoustic analysis; publication dates need not be disturbance dates, and record counts are not independent complaint events or population complaint rates.

**Supplementary Methods S2. Daily detection process**

**S2.1 Separating weekly acoustic opportunity from daily detection**

The 9,912 effort-confirmed station-days described in S1 formed 1,500 station-weeks, each with up to seven daily surveys. Let $z_{it}$ indicate whether Koel acoustic opportunity was present at station $i$ in week $t$, and let $d_{itj} = 1$ when survey day $j$ contained at least one retained Koel-positive sampled minute. We modelled

$$z_{it} \sim \mathrm{Bernoulli}(\psi_{it}), \qquad d_{itj} \mid z_{it} \sim \mathrm{Bernoulli}(z_{it} p_{itj}).$$

Here, $\psi_{it}$ is weekly acoustic opportunity and $p_{itj}$ is the probability that at least one Koel-positive sampled minute is retained on a surveyed day, conditional on that weekly opportunity (*31*). Thus, an all-zero week may indicate either no weekly opportunity or an opportunity that was not observed on any surveyed day. The quantity $p_{itj}$ is an observation probability, not classifier sensitivity or daily abundance. A detection-level station intercept, $u_i \sim \mathrm{Normal}(0, \sigma_p^2)$, represented persistent differences in recorder placement, sound transmission and local recording conditions.

**S2.2 Variables describing daily recording conditions**

Daily observation probability was related to three effort-supported variables. Recording effort was the logarithm of one plus recording hours. Anthropogenic-event rate combined classified car-horn, human-knocking, human-voice and music events per recording hour. Frog-event rate was the corresponding number of classified frog events per recording hour. Both event rates were transformed as $\log(1+x)$ to reduce leverage from strongly right-skewed values. All predictors were standardised before fitting and, during cross-validation, using training stations only.

The retained model was

$$\mathrm{logit}(p_{itj}) = \alpha_0 + \alpha_1 \log\widetilde{(1+R_{itj})} + \alpha_2 \log\widetilde{(1+A_{itj})} + \alpha_3 \log\widetilde{(1+F_{itj})} + u_i,$$

where $R_{itj}$ is recording time in hours and $A_{itj}$ and $F_{itj}$ are anthropogenic- and frog-event rates. These variables were available for every effort-confirmed survey and therefore did not determine which days entered the response. They describe the daily observation environment and should not be interpreted as ecological causes of weekly Koel occurrence.

**S2.3 Selecting the daily detection model**

Candidate models asked four questions: (1) whether effort alone was sufficient; (2) whether short-term acoustic conditions improved observation; (3) whether those conditions should be separated into day-within-week, week-within-station and between-station components; and (4) whether frog-event rate required transformation or the near-constant insect-event rate added useful information. All candidates used the same effort-defined response, weekly occurrence structure, priors, detection-level station intercept and five whole-station folds.

Complete held-station detection-history log predictive density was the primary comparison. Models within one site-clustered standard error of the numerical leader formed the predictive-equivalence set; within that set, we retained the model with the fewest detection terms. The separated within-between model had the highest numerical predictive density, but the three-term model above remained within this equivalence set. We preferred the transformed frog rate because its raw distribution had a strong right tail. We removed insect-event rate because it contained little variation and improved predictive density by only $1.37 \pm 1.74$ units when added to the otherwise identical model. Candidate definitions and validation results are reported in Supplementary Data 4.

**S2.4 Checking dependence and sound-pressure bias**

Each candidate first required complete finite inputs, non-zero information and a full-rank design. Within the detection layer, $|r| = 0.70$ and variance-inflation factor (VIF) = 5 triggered review; $|r| = 0.85$ or VIF = 10 indicated severe dependence (*52*). The retained design was complete and full rank, with maximum $|r| = 0.097$ and VIF = 1.021.

Detection and occurrence predictors enter different likelihood components, so we did not calculate one combined VIF. Instead, daily observation variables were aggregated to station-weeks and compared with occurrence predictors using Pearson and Spearman associations. The largest absolute association was 0.465, between frog-event rate and the annual cosine term, below the 0.70 review threshold. This check found no strong pairwise cross-layer dependence; it does not prove complete separation of ecological and observation processes.

Event-sampled 0-2-kHz sound pressure level was unavailable on days without a retained bird event. A second version excluding Koel-positive sampled minutes remained response-derived. The two versions were therefore evaluated only as paired diagnostics on 9,561 days and 1,471 station-weeks. Neither improved diagnostic predictive density relative to the matched model without sound pressure. They were not treated as all-minute ambient sound or used to select the detection model. Supplementary Data 5 records these design, dependence and response-dependence checks.

**S2.5 Whole-station validation and use in the joint model**

Fivefold cross-validation withheld complete monitoring stations. Each training fold supplied predictor means and standard deviations, and the detection-level station effect at a withheld station was integrated over its estimated distribution. We scored the complete held-station daily history. Predictive density determined the equivalence set; Brier score, calibration, predicted detection counts, all-zero histories and MCMC diagnostics provided supporting checks. Fig. S2 presents these out-of-fold diagnostics for the retained model.

The retained formula was confirmed after refitting the ecological models and was then kept unchanged. It enters the final joint model unchanged. S5 adds the positive-minute count model for days with detections; it does not replace the imperfect-detection component defined here.

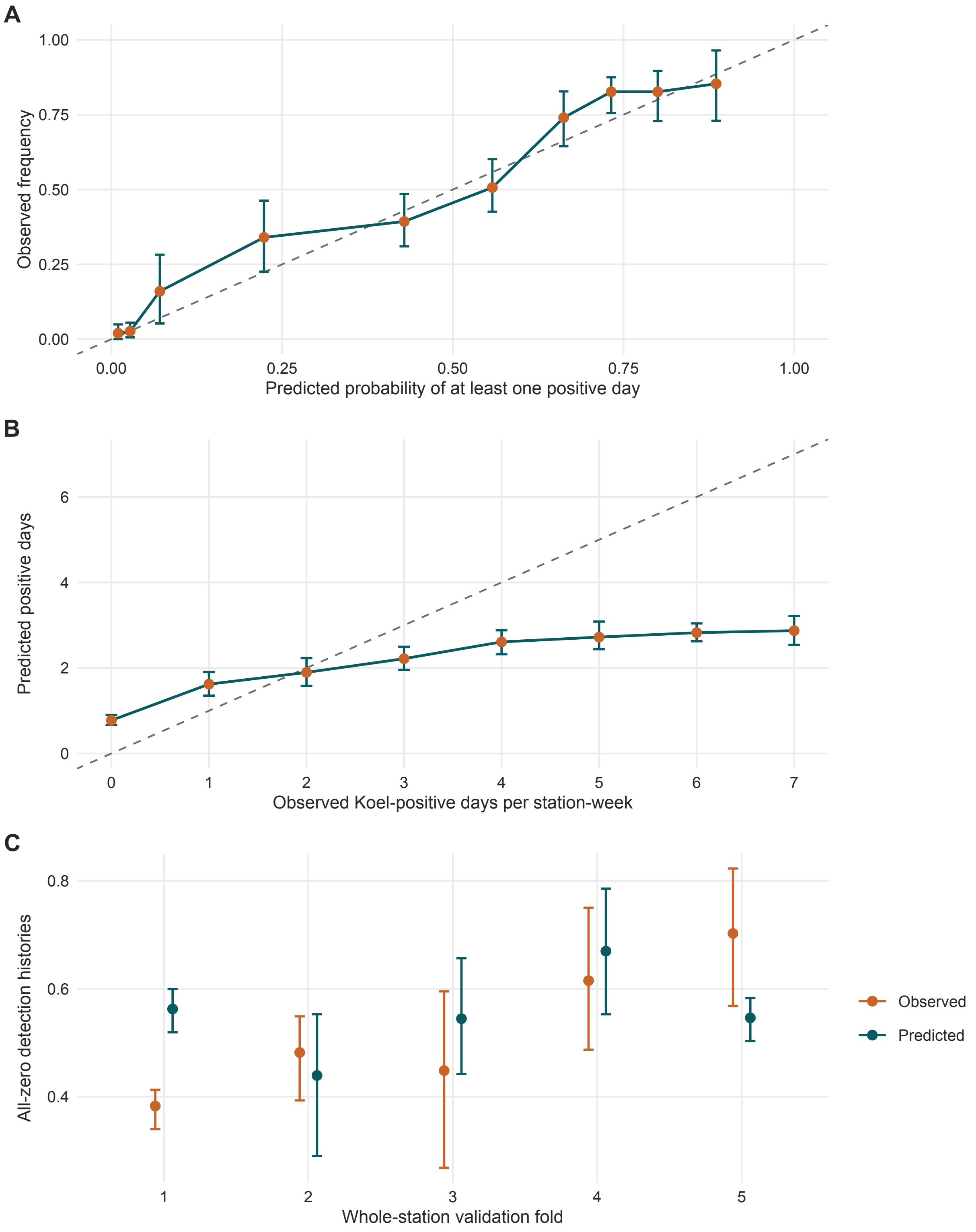


Fig. S2. Whole-station validation of the retained daily-detection model. (A) Calibration of the predicted probability of at least one Koel-positive day, summarized across ten equally sized prediction groups. (B) Mean predicted number of positive days within each observed-count group. (C) Observed and predicted proportions of all-zero daily histories within each held-station fold. All predictions were generated out of fold for 31 stations and 1,500 station-weeks. Error bars are 95% monitoring-station-cluster bootstrap intervals, stratified by validation fold; dashed grey lines indicate perfect agreement.

### Supplementary Methods S3. Environmental variables and potential-host models

#### S3.1 Spatial units used for fitting and city prediction

Environmental variables were calculated around each recorder. Coordinates were transformed from WGS 84 (EPSG:4326) to China Geodetic Coordinate System 2000 / 3-degree Gauss–Krüger CM 114° E (EPSG:4547) before metric operations. The main analysis used station-centred 250-m-radius circles; matching 100- and 500-m circles supported the scale sensitivity in S4.

For city prediction, the same summaries were recalculated within 250-m circles around fixed hexagonal reporting-cell centroids (hereafter, HEX centroids). HEX polygons stored and displayed predictions but did not define the ecological neighbourhood. Administrative outlines were cartographic only.

#### S3.2 Environmental data and extraction

Land-cover variables came from Dynamic World V1 in Google Earth Engine (*41*, *47*). We averaged its class-probability bands from 1 March 2023 to 1 April 2026 and summarized the required bands within each circle. Retained models used tree, built and grass probabilities. These soft probabilities are not mutually exclusive hard land-cover fractions.

Tree-edge area used the same composite. Pixels with mean tree probability at least 0.50 were classified as tree pixels; local tree/non-tree boundaries were identified from 50-m focal maximum and minimum filters, and boundary-pixel area was summed within each circle. This is a structural proxy, not a mapped biological forest edge.

Distance to recurrent water used the 30-m JRC Global Surface Water v1.4 occurrence layer (*43*). Pixels with occurrence at least 50% defined recurrent water. Euclidean distance was sampled at each station or HEX centroid, not minimized within its circle. Mean elevation came from NASA SRTMGL1 v3 (*42*). ESA WorldCover 2021 v200 class 80 supplied a separate permanent-water reporting check and did not enter the occurrence formulae (*46*).

Major-road density was derived from a dated OpenStreetMap snapshot. Selected major-road centre-lines were clipped to each 250-m circular neighbourhood, and density was calculated as the total road length within the neighbourhood per unit area (km of road per $km^2$). Supplementary Data 6 defines all analytical variables and their interpretation limits, while Supplementary Data 7 reports data sources, licences and processing steps.

Continuous predictors were standardized before fitting. Whole-station validation estimated means and standard deviations from training stations only. Missing or non-finite required values made a station-week or HEX-week invalid rather than zero.

#### S3.3 Defining the three potential-host acoustic responses

Potential-host responses used the same effort-confirmed days as the Koel response. An assemblage day was positive when at least one member had a retained acoustic record.

1. The Corvid assemblage comprised Red-billed Blue Magpie (*Urocissa erythrorhyncha*),

Azure-winged Magpie (*Cyanopica cyanus*), Oriental Magpie (*Pica serica*) and Large-billed Crow (*Corvus macrorhynchos*).

2. The Sturnid assemblage comprised Black-collared Starling (*Gracupica nigricollis*), Common Myna (*Acridotheres tristis*) and Crested Myna (*Acridotheres cristatellus*).

3. Masked Laughingthrush (*Pterorhinus perspicillatus*) was modelled as a separate response.

These response definitions were fixed before candidate models were compared. They represent potential acoustic co-occurrence with Koel. They do not establish brood-host use, host preference, abundance or causation. Species membership and daily response rules are listed in Supplementary Data 8 (Table S8A).

**S3.4 Selecting one model for each potential-host response**

Each response used the weekly-state model in S2 and the same daily-detection formula, which was kept unchanged:

$$\operatorname{logit}(p_{itj}) = \alpha_0 + \alpha_1 \log\big(\widetilde{1 + R_{itj}}\big) + \alpha_2 \log\big(\widetilde{1 + A_{itj}}\big) + \alpha_3 \log\big(\widetilde{1 + F_{itj}}\big) + u_i,$$

where $R$ is recording time, $A$ is anthropogenic-event rate, $F$ is frog-event rate and $u_i$ is the detection-level station intercept. Host-model development then followed six steps.

1. Define small candidate sets. Candidate variables were grouped by ecological role rather than combined freely. Corvid models contrasted tree cover or tree-edge structure

with built cover or major-road density. Sturnid models contrasted grass or open-green cover with built cover or major-road density. Masked Laughingthrush models contrasted shrub cover or tree-edge structure with built cover. Distance to water, elevation and the annual sine and cosine terms completed the candidate sets. The full formulae are listed in Supplementary Data 8 (Table S8B).

2. Check data completeness and model design. A candidate proceeded only when all required values were finite for the effort-confirmed surveys included in the analysis, the design matrix had full rank, and no predictor had near-zero variance. Near-zero variance meant a standard deviation below $10^{-6}$ or fewer than three unique finite values.

3. Review collinearity. We flagged a candidate when the largest absolute pairwise correlation was at least 0.70 or the largest variance inflation factor (VIF) was at least 5. Values of $|r| \geq 0.85$ or VIF $\geq 10$ were treated as severe. A severe flag prevented independent coefficient interpretation (*52*). A prediction-only candidate could proceed only when its design matrix remained full rank and whole-station validation showed stable transfer; correlated coefficients were not interpreted separately.

4. Check Bayesian sampling. Short fits tested the model matrix and sampler before formal fitting. These checks did not rank candidates. Formal fits were classified as clean when $\hat{R} \leq 1.05$ and effective sample size was at least 400, acceptable when $\hat{R} \leq 1.10$ and effective sample size was at least 200, and otherwise of concern (*53*).

5. Compare candidate structures. Stable formal fits were ranked by full-data PSIS-LOO predictive density for the complete daily detection history (*54*). Paired differences were aggregated to the 31 stations. The numerical leader was retained provisionally, while candidates within one station-clustered standard error were treated as statistically unresolved. Coefficient significance was not used to delete variables.

6. Select one formula for subsequent analyses. The retained models were C02 for Corvids, S03 for Sturnids and L02 for Masked Laughingthrush:

$$\begin{aligned}\text{Corvid opportunity} &\sim \text{trees} + \text{built} + \text{distance to water}\\ &\quad +\text{elevation} + \sin(\text{year}) + \cos(\text{year})\,,\\ \text{Sturnid opportunity} &\sim \text{grass} + \text{major-road density} + \text{distance to water}\\ &\quad +\text{elevation} + \sin(\text{year}) + \cos(\text{year})\,,\\ \text{Masked Laughingthrush opportunity} &\sim \text{tree-edge area} + \text{built} + \text{distance to water}\\ &\quad +\text{elevation} + \sin(\text{year}) + \cos(\text{year})\,.\end{aligned}$$

C02 and L02 had the highest predictive scores within their one-standard-error sets and were retained for subsequent analyses, not as uniquely supported ecological explanations. The sparser Sturnid response required a direct whole-station comparison of S01–S03. S03 had the highest held-station predictive density, but all pairwise 95% intervals included zero and S02 had better Brier score and calibration. S03 was therefore retained for subsequent analyses with this uncertainty stated explicitly.

The three selected models were then evaluated with the same five whole-station folds. Scaling and fitting used training stations only, and complete daily histories were scored at withheld stations. Expected log predictive density (ELPD) was primary; Brier score and calibration were supporting diagnostics. Calibration slopes were 0.755 for Corvids, 0.246 for Sturnids and 0.758 for Masked Laughingthrush. The weak Sturnid calibration was reported explicitly; posterior uncertainty from this model was propagated into Koel inference. Supplementary Data 8 (Table S8C) reports the validation results and fig. S3 shows out-of-fold calibration.

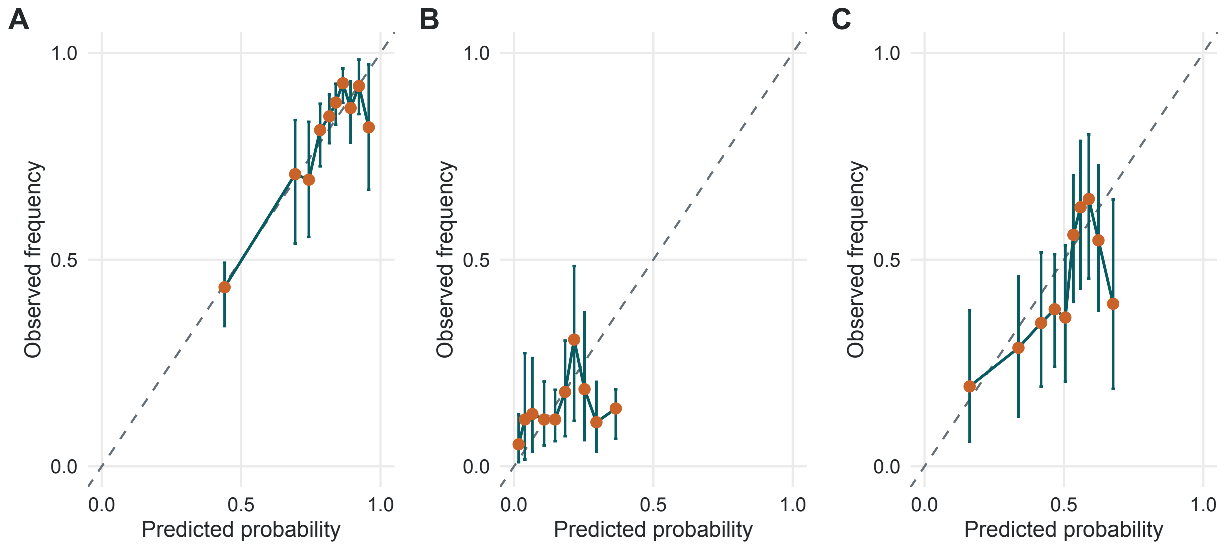


Fig. S3. Whole-station calibration of the three potential-host acoustic-opportunity models. Points show observed station-week frequencies within ten equally sized groups of out-of-fold predicted probabilities for (A) the Corvid assemblage, (B) the Sturnid assemblage and (C) Masked Laughingthrush. The dashed line denotes perfect calibration, and vertical bars show 95% monitoring-station-cluster bootstrap intervals for the observed frequency. Calibration compared the predicted probability of at least one recorded positive day within a station-week with the observed weekly detection outcome. All predictions were generated for stations excluded from fitting. These responses quantify potential acoustic co-occurrence and do not establish brood-host use.

#### S3.5 Passing host uncertainty into the Koel model

Host opportunity was not treated as a known covariate. Within each validation fold, uncertainty was passed forward in four steps.

1. Fit each selected host model using training stations only.

2. Draw weekly host opportunity for both training and withheld stations from each host posterior.

3. Standardise each host draw using that host model's training-fold mean and standard deviation.

4. Fit the Koel model under three independently sampled, complete host-opportunity surfaces and pool equal numbers of the resulting posterior draws.

This procedure carried uncertainty from all three host models into Koel predictions while keeping test stations outside fitting and standardisation. Three posterior surfaces were a validation-level computational compromise: they allowed transfer scores to reflect host uncertainty but did not demonstrate that coefficient credible intervals had stabilised with respect to the number of sampled surfaces.

After validation, the selected host models were refitted using all 31 stations and projected to city HEX centroids with the same 250-m predictors. Their standardised posterior opportunities entered the Koel model as one joint predictive block. Individual coefficients from this correlated block were not interpreted as independent host importance. An equal-weight index, calculated after separately standardising the three response groups, was used only as a compression sensitivity and in the calling-density branch. Equal weight refers to response groups, not species abundance or biological importance.

**Supplementary Methods S4. Koel acoustic opportunity, model development and spatial transfer**

**S4.1 Order of model development**

This section concerns weekly Koel acoustic opportunity, $\psi_{it}$. The daily-detection model in S2 was kept unchanged, and the positive-minute component is introduced in S5. Koel model development followed a fixed order:

1. finalise the corrected daily response and detection model (S1–S2);

2. select one occurrence model for each potential-host response and keep its formula unchanged (S3);

3. select a Koel landscape-season base model without host terms;

4. add the predefined host structures H0–H5;

5. fix all transfer formulae before whole-station cross-validation; and

6. compare spatial scale, coordinate trends and residual spatial pattern without returning to variable selection.

This order kept observation-process decisions, host estimation, Koel structure development and transfer testing separate. Supplementary Data 9 (Table S9A) records the input and output of each stage.

#### S4.2 Selecting the Koel landscape-season base model

The Koel landscape candidate pool contained tree cover, tree-edge area, built cover, open-green cover, major-road density, distance to recurrent water and elevation. All candidates also contained the annual sine and cosine terms. Four small formulae were fixed before formal fitting:

$K01$:tree edge + built + distance to water + elevation + annual cycle,
$K02$:trees + built + distance to water + elevation + annual cycle,
$K03$:tree edge + major-road density + distance to water + elevation + annual cycle,
$K04$:tree edge + open green + built + elevation + annual cycle.

The same checks of completeness, variance, rank and collinearity used for the host models were applied before fitting. Short fits checked sampling, and stable formal fits were compared by full-data PSIS-LOO density for the complete detection history (*54*). The candidate with the highest stable ELPD was retained, while candidates within one station-clustered standard error were treated as unresolved. Credible intervals for individual coefficients were not used as a deletion rule.

K02 had the highest ELPD (–3537.24). K03 was 6.31 units lower with a station-clustered standard error of 7.91 and therefore remained in the one-standard-error set. K02 was retained as the base model for subsequent analyses. Its largest predictor correlation was 0.816 and its largest VIF was 3.71. This exceeded the correlation-review threshold but not the threshold for severe dependence; tree and built cover were retained as distinct landscape descriptors. Supplementary Data 9 (Table S9B) reports all four candidates and the model-selection decision.

**S4.3 Adding potential-host opportunity and defining model roles**

Keeping the K02 base model unchanged, we defined six host structures:

1. H0: no host term;

2. H1: Corvid opportunity only;

3. H2: Sturnid opportunity only;

4. H3: Masked Laughingthrush opportunity only;

5. H4: the three host opportunities entered separately as one predictive block; and

6. H5: one equal-weight index formed after separately standardising the three host opportunities.

The models then separated into two branches.

Monitored-site association branch. H1, H2 and H3 were each fitted with an occurrence-level station intercept, $b_i$:

$$\text{logit}(\psi_{it}) = \beta_0 + \mathbf{x}_{it}^{\top}\boldsymbol{\beta} + \beta_H H_{it} + b_i.$$

These models asked whether Koel and one potential-host opportunity covaried after accounting for the common landscape-season predictors and persistent differences among the monitored communities. They were parallel association analyses, not steps for deciding which host to retain. Their coefficients do not establish host use or causation (fig. S4A).

Unmonitored-site transfer branch. H0, H4 and H5 omitted the occurrence-level station intercept because a station-specific value cannot be assigned to an unmonitored location. Each structure was fitted without coordinates (P0) and with standardised projected easting and northing (Pxy). Coordinates represented broad residual geography, not ecological mechanisms. The detection-level station intercept remained in every model and was integrated over its estimated distribution for a withheld station.

H4 was evaluated as a joint predictive host block. Its correlated host coefficients were not used to rank hosts and were not interpreted as independent biological effects. H5 tested whether the same information could be compressed into one equal-scale index. The complete list of models and their interpretation limits are given in Supplementary Data 9 (Table S9C).

Host uncertainty was propagated in every H1–H5 fit. Three independent posterior draws from each selected host model were standardised within the relevant training data and passed into separate Koel fits. Posterior draws from the three fits were pooled with equal weights. Pairwise Spearman correlations among the three H4 $\psi$ surfaces were 0.992, indicating closely matching spatial rankings across imputations.

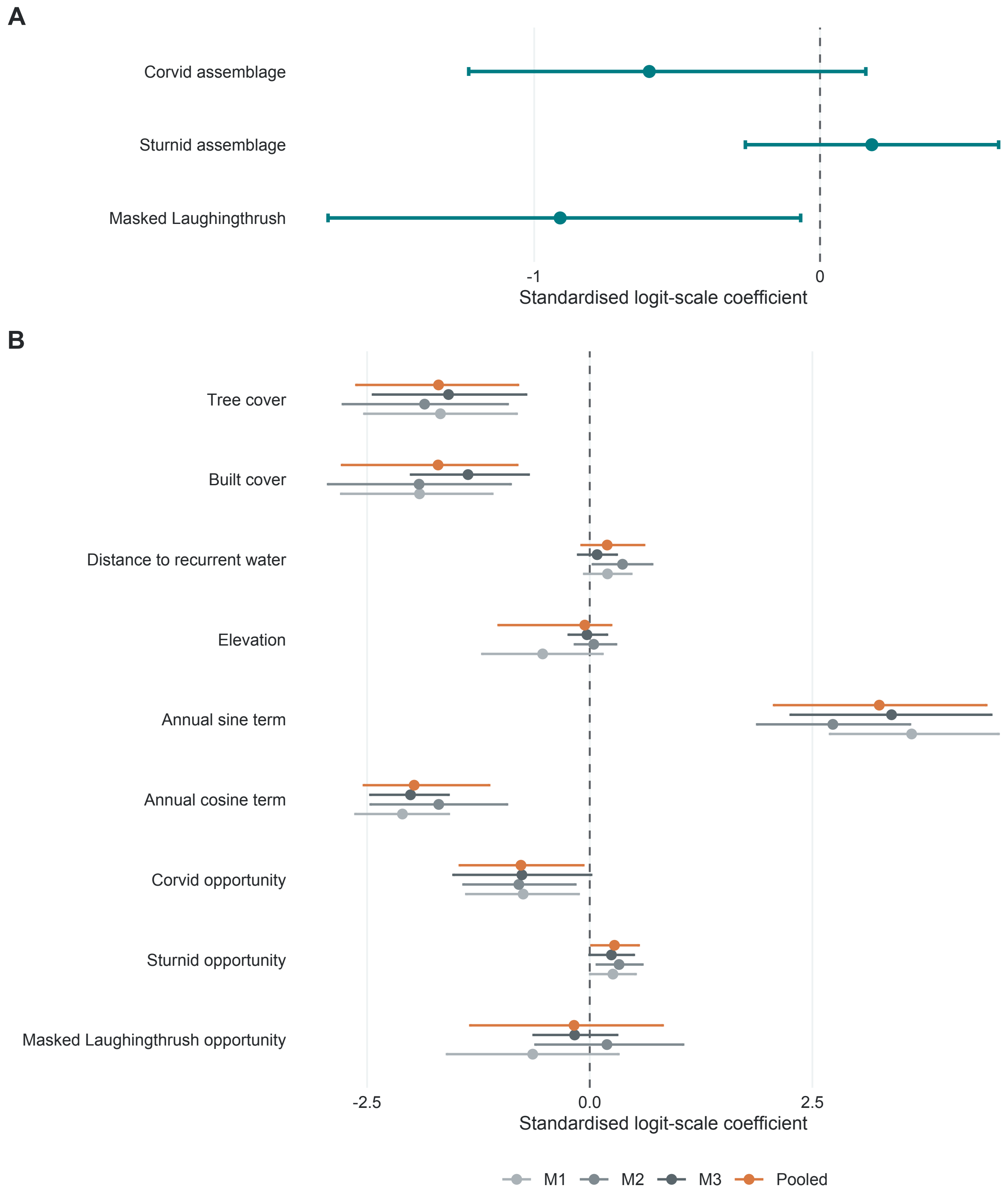


Fig. S4. Potential-host associations and propagation of host-model uncertainty into Koel weekly acoustic opportunity. (A) Posterior median and 95% credible interval for the host term in each monitored-site single-host model. Each model contained the K02 landscape-season predictors and an occurrence-level station intercept. (B) K02 and host coefficients from H4 P0 under three independently sampled host-opportunity surfaces (M1–M3) and their equal-weight pooled posterior. This panel shows propagation across imputations; it does not compare propagated estimates with an error-free plug-in fit. Single-host terms describe conditional acoustic association, while the H4 host terms form one predictive block. Neither establishes brood-host use or causation.

**S4.4 Testing transfer to unmonitored stations**

All transfer formulae were fixed before fivefold whole-station cross-validation. Each fold followed the same sequence:

1. withhold every day and week from the test stations;

2. calculate predictor means and standard deviations from training stations only;

3. refit the three host models and predict host opportunity for training and test stations;

4. fit each prespecified Koel structure under the three sampled host-opportunity surfaces and pool the resulting posterior predictions; and

5. score the complete daily detection histories at the withheld stations.

Joint held-station ELPD was the primary score. Brier score for at least one weekly detection, calibration intercept and slope, fold influence, posterior diagnostics and residual spatial autocorrelation were supporting checks. All 85 expected fits met the diagnostic criteria (maximum $\hat{R} = 1.040$; minimum bulk effective sample size = 210.6).

Because candidate formulae were not reselected within each fold, the results measure transfer performance conditional on the selected model structures. They do not represent nested model-selection cross-validation. Supplementary Data 10 reports model-level metrics and paired station-clustered contrasts.

#### S4.5 Selecting the host-informed planning formulation

The primary host-informed ecological-planning formulation was H4 P0 at 250 m:

$$\begin{aligned}\operatorname{logit}(\psi_{it}) = \beta_0 + \beta_1\,\text{trees}_i + \beta_2\,\text{built}_i + \beta_3\,\text{distance to water}_i + \beta_4\,\text{elevation}_i \\ +\beta_5 \sin(\text{year}_t) + \beta_6 \cos(\text{year}_t) + \beta_7 H_{it}^{\text{corvid}} + \beta_8 H_{it}^{\text{sturnid}} + \beta_9 H_{it}^{\text{laughingthrush}}.\end{aligned}$$

H4 P0 improved held-station ELPD by 17.96 units relative to the matched coordinate-free no-host model (95% site-clustered interval 0.41–35.51). Adding coordinates to H4 changed ELPD by 15.39 units, but its interval included zero (–3.18–33.97). H4 P0 also showed no detectable residual spatial autocorrelation. These results supported H4 P0 as the main host-informed model without adding a non-mechanistic coordinate trend.

This choice is question-specific, not a claim that H4 P0 had the highest ELPD among every candidate. Under the stricter all-model transfer rule, H0 Pxy was the conservative numerical benchmark; H4 Pxy differed from it by only 0.79 ELPD units (–5.27–6.85). H0 Pxy, H4 Pxy and H5 therefore remain transparent transfer sensitivities. Citywide host-informed interpretation uses H4 P0, while conclusions that depend on the coordinate benchmark must be reported as sensitivity results.

#### S4.6 Comparing 100-, 250- and 500-m neighbourhoods

The scale comparison used the same response, folds, detection model, priors and training-fold standardisation at all three radii (*44*). A matched no-host Pxy structure was used so that the test isolated the landscape neighbourhood rather than changes in the host block.

Relative to 250 m, paired ELPD was −5.93 units at 100 m (95% interval −13.37–1.52) and −1.41 units at 500 m (−7.24–4.41). Neither alternative clearly improved transfer. We therefore used 250 m for subsequent analyses and evaluated H4 at that scale. This result does not identify 250 m as a unique biological scale.

#### S4.7 Checking residual spatial autocorrelation

Residual spatial structure was checked separately for each of the six 250-m transfer models. Out-of-fold detection-history Pearson residuals were aggregated to the 31 stations. We used a symmetric four-nearest-neighbour graph with inverse-distance, row-standardised weights. Moran's $I$ was tested with 9,999 permutations, and probabilities were adjusted across the six models with the Benjamini–Hochberg method (*55*).

H4 P0 had Moran's $I = 0.025$ (two-sided permutation $p = 0.586$, adjusted $p = 0.987$). No tested structure retained detectable residual spatial autocorrelation. This diagnostic supported the absence of a remaining spatial pattern in the held-station residuals; it did not replace held-station predictive scoring or prove that all spatial processes had been measured. Fig. S5 summarises the scale, model and residual-spatial comparisons.

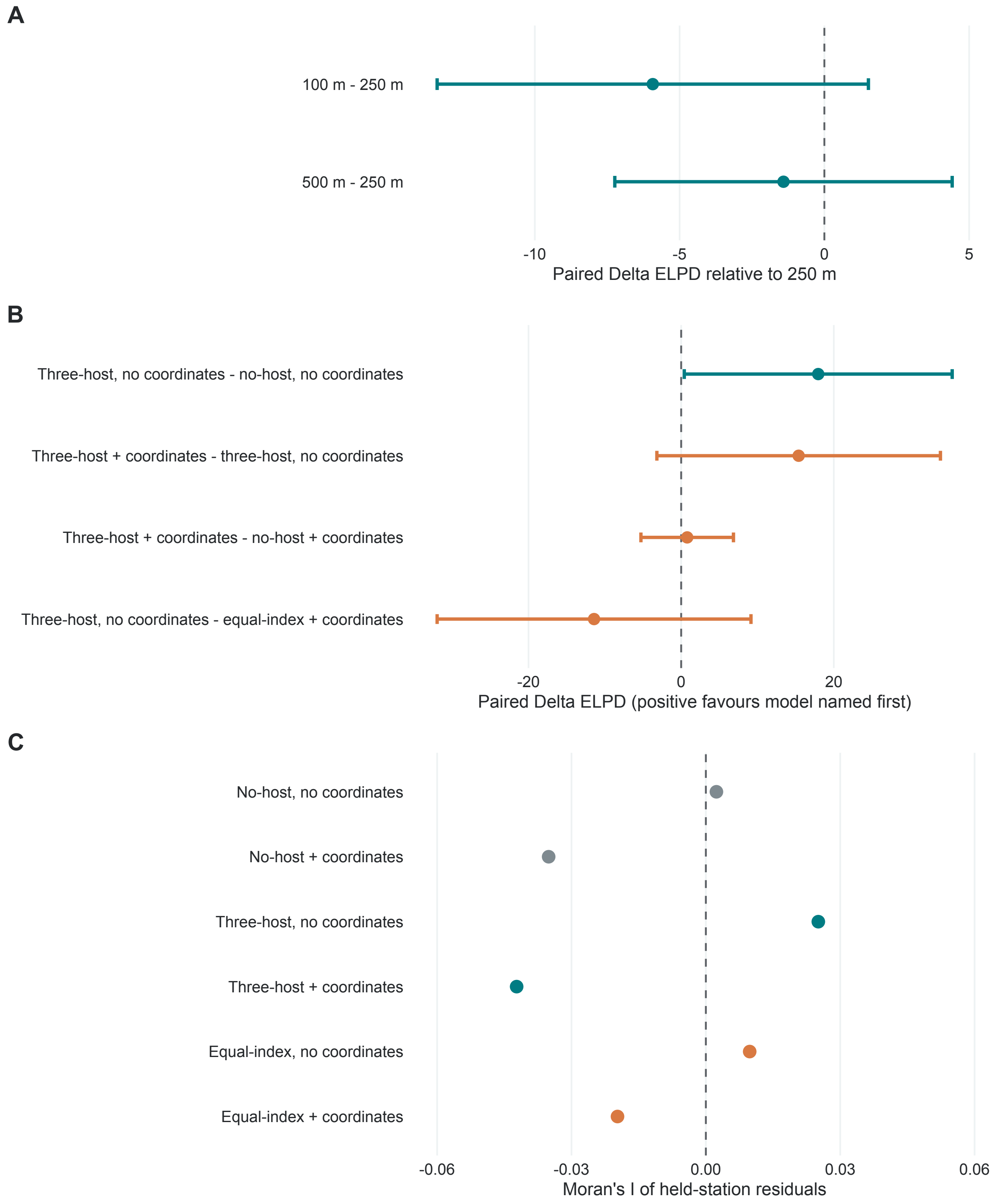


Fig. S5. Whole-station transfer, scale and residual-spatial diagnostics for Koel weekly acoustic opportunity. (A) Paired change in held-station ELPD for 100- and 500-m landscape neighbourhoods relative to 250 m under the matched no-host coordinate-trend structure. (B) Prespecified 250-m contrasts testing the joint host block, coordinate trend and equal-index sensitivity; positive values favour the model named first. In panels A and B, points show paired ELPD differences, and horizontal bars show 95% station-clustered intervals. Panels A and B come from separate matched validation stages, so absolute ELPD values should not be compared across panels. (C) Moran's I for out-of-fold station-level detection-history residuals from the six transfer models. Benjamini–Hochberg-adjusted permutation probabilities are reported in Supplementary Data 10 (Table S10B). Formulae were fixed before whole-station validation.

### Supplementary Methods S5. Joint calling density and sampled-minute acoustic exposure

#### S5.1 Linking weekly opportunity, daily detection and positive-minute counts

The weekly-state model in S4 and the daily-detection model in S2 were extended with a third component for the number of Koel-positive sampled minutes. For station $i$, week $t$ and surveyed day $j$, let $K_{itj}$ be the number of Koel-positive sampled minutes among $N_{itj}$ recorded minutes, and let $d_{itj} = 1(K_{itj} > 0)$. The daily joint model was

$$z_{it} \sim \text{Bernoulli}(\psi_{it}), \qquad d_{itj} \mid z_{it} \sim \text{Bernoulli}(z_{it} p_{itj}),$$

$$K_{itj} = 0 \quad \text{if } d_{itj} = 0, \qquad K_{itj} \mid d_{itj} = 1 \sim \text{ZT-BetaBinomial}(N_{itj}, \mu_{it}, \phi).$$

Here, $\psi_{it}$ is weekly acoustic opportunity, $p_{itj}$ is the probability of retaining at least one Koel-positive minute on a surveyed day when that opportunity is present, and $\mu_{it}$ controls positive-minute density. The zero-truncated count distribution is conditional on a positive survey day. Its mean parameter is therefore not an unconditional minute-level probability.

All three components were estimated in one daily likelihood. All surveyed days informed weekly opportunity and daily detectability, while the positive-minute counts additionally informed calling density. This construction retained imperfect detection rather than replacing the occupancy model with a count model. It also prevented missing days from being treated as zero counts. Supplementary Data 11 (Table S11A) lists the three likelihood components and their roles.

### S5.2 Modelling host-associated daily calling density

We first compared two daily calling-density structures. The seasonal model contained annual sine and cosine terms, whereas the host-informed model added one equal-scale potential-host index. During fitting, annual seasonality used the dated sine and cosine terms defined in S1.2. City predictions were evaluated on a common 52-week annual prototype:

$$\mathrm{logit}(\mu_{it}) = \gamma_0 + \gamma_H H_i^{\mathrm{eq}} + \gamma_1 \sin_t + \gamma_2 \cos_t.$$

For reporting, W01–W52 formed a common annual prototype beginning on 1 January. Observation dates were assigned to seven-day day-of-year bins, with the final bin capped at W52. This reporting axis was separate from the dated seasonal basis used for fitting. The index $H_i^{\mathrm{eq}}$ was constructed in four steps:

1. For each potential-host response in S3, we predicted mean weekly acoustic opportunity across a common 365-day annual clock.

2. We standardised the Corvid, Sturnid and Masked Laughingthrush predictions separately.

3. We averaged the three standardised response-group values with equal weight.

4. We standardised the resulting index again before fitting.

Full-data scaling used all 31 stations. Each validation fold used training stations only. Equal weighting placed the three response groups on a common scale; it did not weight species by record count, abundance or presumed biological importance.

We fitted the density model under three independently sampled host-opportunity surfaces and pooled equal numbers of posterior draws. The prespecified HEQZ formulation was retained as the host-informed ecological and planning branch because its host coefficient was positive and the overall held-station contrast favoured HEQZ; the season-only S1 model remained a sensitivity because the gain was not consistent across all five folds (Supplementary Data 11 (Table S11C)). Coefficient posteriors are shown in fig. S6A; the host coefficient and model-validation summaries are reported in Supplementary Data 11 (Table S11C).

The host term describes association between long-term potential-host acoustic opportunity and Koel-positive minute density. It is not evidence of host use, parasitism, abundance or a causal effect on Koel calling.

#### S5.3 Resolving calling density by hour

The daily count model described annual change but not the distribution of calling within a day. We therefore assembled hourly records from the same 9,912 effort-confirmed survey days. For each recorded clock hour $h$, $N_{itjh}$ was the number of sampled minutes and $K_{itjh}$ was the number classified as Koel-positive. Hours without recording effort remained missing. Within the 2,682 positive survey days, all 64,141 recorded hour cells were retained, including 56,348 cells with zero Koel-positive minutes.

The hourly density screen conditioned on a positive survey day and fitted all recorded hours together:

$$K_{itjh} \sim \mathrm{BetaBinomial}\big(N_{itjh}, \mu_{itjh}, \phi\big), \quad \text{conditional on } \sum_h K_{itjh} > 0,$$

$$\mathrm{logit}\big(\mu_{itjh}\big) = \mathbf{x}_{th}^{\top}\boldsymbol{\gamma} + v_{itj}, \qquad v_{itj} \sim \mathcal{N}\big(0, \sigma_{\mathrm{day}}^2\big).$$

The day intercept $v_{itj}$ represented residual variation shared among hours of the same survey day. Three temporal structures were specified before comparison.

1. D0 contained the annual sine and cosine terms.

2. D1 added two cyclic Fourier harmonics for clock hour.

3. D2 added interactions between annual phase and the four hourly terms.

Candidate screening used a fixed 600-day subset spanning all stations, both years and the five validation folds. Transfer was then tested by withholding complete stations, fitting four folds and scoring positive-day hour histories at the omitted stations. These validation fits used posterior modes under the same priors and integrated the new-day intercept by Gaussian quadrature. The retained structure was subsequently fitted to all positive days with four MCMC chains.

D1 clearly improved held-station predictive density over D0. D2 provided only a small and uncertain improvement over D1, so D1 was retained and D2 became a sensitivity. The D1 posterior reproduced the mean and upper tail of daily positive-minute counts, the fraction of recorded hours with zero counts, and the observed 04:00-08:00 and 15:00-19:00 shares. Candidate definitions and diagnostics are reported in Supplementary Data 11 (Table S11B-C).

#### S5.4 Combining the daily and hourly components

After selecting D1, we replaced the daily density branch with the hourly branch in one confirmatory integrated fit while leaving the H4 P0 occurrence formula, daily-detection formula, host inputs and priors unchanged. This fit tested whether the added hourly information altered the common $\psi$- and $p$-parameters. The largest shift was 0.057 standard deviations of the earlier posterior, and the integrated fit met the MCMC diagnostic criteria.

The final host-informed hourly prediction therefore used a cut-posterior composition of completed models. The daily host-informed fit supplied the intercept, equal-scale host effect and annual terms. D1 supplied the two clock-hour harmonics, positive-day dispersion and day-level variation. Day-level variation was propagated through posterior prediction. The urban-overlap summaries used a standard detection reference with the detection-site intercept fixed at zero. Only completed posterior draws entered prediction; an incomplete direct host-by-hour refit did not contribute estimates.

This modular step did not change the selected occurrence, detection or host formulae. It added hourly resolution after checking that the hourly branch did not materially change the shared parameters. Supplementary Data 11 distinguishes the fully joint daily fit, the modular hourly screen and the confirmatory integrated fit.

**S5.5 Derived sampled-minute metrics**

The hourly count model was fitted conditional on a positive day. We therefore denote its beta-binomial mean by $\mu$, and the corresponding zero-truncation-adjusted probability that a sampled minute in hour h was Koel-positive by $\mu^{+}$:

$$\mu^{+}_{itjh} = \frac{\mu_{itjh}}{1 - P\left(\sum_h K_{itjh} = 0\right)}.$$

Posterior draws from the three components were then combined before any summary was calculated:

$$E_{itjh} = \psi_{it}\, p_{itj}\, \mu^{+}_{itjh}.$$

The resulting $E_{itjh}$ is the model-implied probability that a sampled minute, under the stated recording and detection reference, is classified as Koel-positive. Expected positive sampled minutes in a time window $W$ were obtained by summing the draw-wise probabilities across its scheduled samples:

$$\mathbb{E}[M_{iW}] = \sum_{(t,j,h)\in W} n^{(W)}_{itjh}\, E_{itjh}.$$

Window-level probabilities were calculated for each posterior draw from the beta-binomial zero-count probabilities, adjusted for conditioning on a positive day, and combined with weekly opportunity and daily detectability. Day-level variation was integrated across the posterior calculation. No independent-minute approximation was used. The standard detection reference used 144 scheduled sampled minutes per day and centred anthropogenic- and frog-event covariates. Annual monitored-site summaries retained observed recording conditions. All-day exposure used the mean of the 24 hourly values because the reference design scheduled six sampled minutes in every hour. Supplementary Data 12 defines each public-facing metric, its denominator and its interpretation boundary.

These quantities describe algorithm-positive sampled minutes under the study design. They are not continuous calling duration, sound-pressure level, perceived loudness, annoyance, abundance, residence, nesting or the probability that a member of the public hears a Koel. The annual and hourly patterns, together with their temporal support, are summarised in fig. S6.

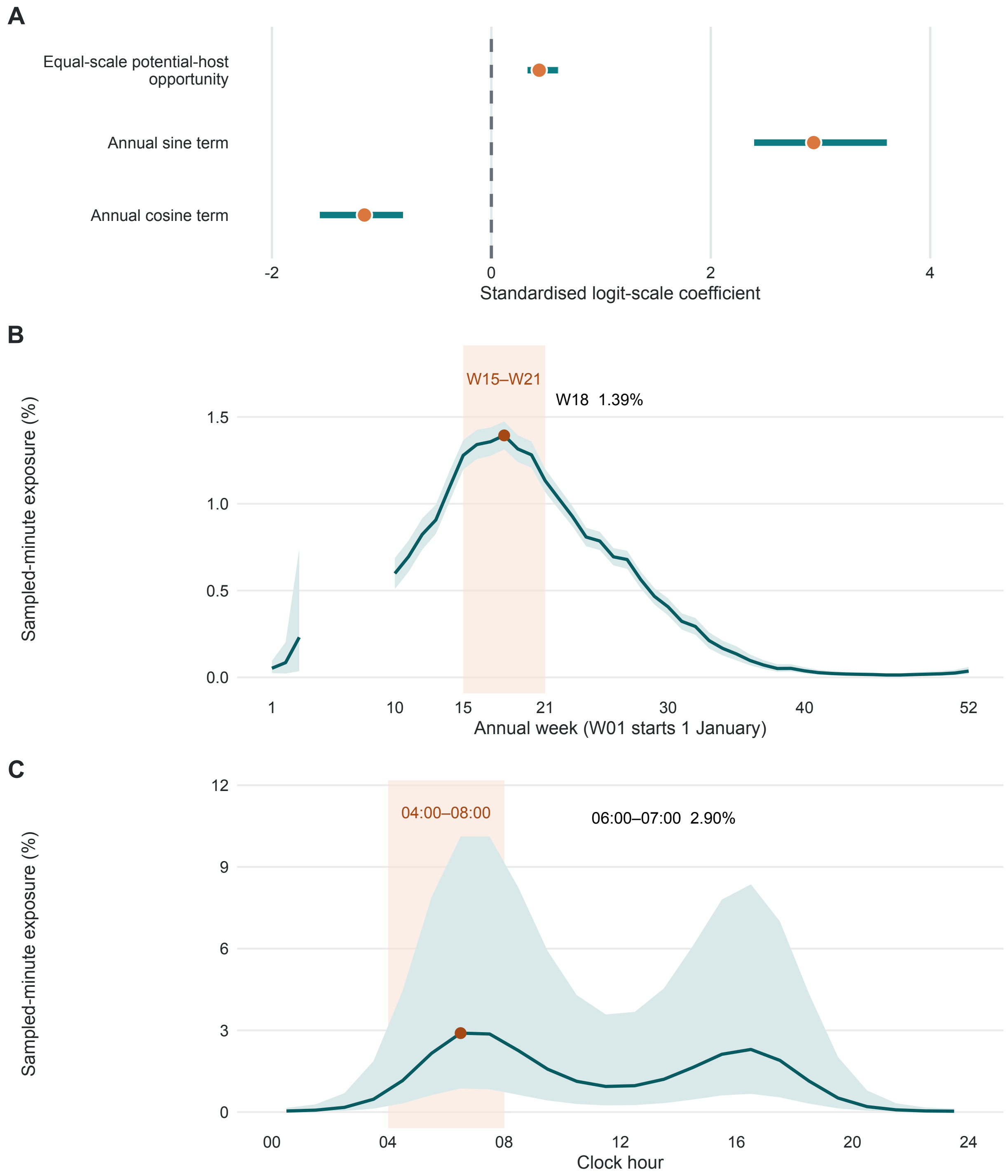


Fig. S6. Host-informed calling density and temporal pattern of sampled-minute acoustic exposure. The fitted calling-density parameter is denoted $\mu$. Its zero-truncation-adjusted positive-day sampled-minute probability, $\mu^+$, was combined with weekly acoustic opportunity $\psi$ and daily detectability $p$ to derive sampled-minute acoustic exposure, $E = \psi p \mu^+$. (A) Posterior medians and 95% credible intervals for the equal-scale potential-host and annual terms in the host-informed daily calling-density branch; the host coefficient and model-validation summaries are reported in Supplementary Data 11 (Table S11C). (B) Annual pattern of monitored-site sampled-minute acoustic exposure under observed recording conditions. Gaps indicate weeks without observations; the pale-orange band marks W15–W21, and the orange point marks the W18 maximum. (C) Hourly sampled-minute acoustic exposure in week 18 under the standard detection reference; the pale-orange band marks 04:00–08:00. Curves show posterior means, and ribbons show 95% credible intervals. Exposure represents the posterior probability that a scheduled sampled minute is classified as Koel-positive under the stated observation conditions; it does not represent sound level, continuous calling duration or realised human disturbance.

### Supplementary Methods S6. City prediction and interpretation support

#### S6.1 Projecting the fixed model across the city

The final city projection used the coordinate-free H4 P0 model retained in S4. The occurrence-level station intercept was omitted because it cannot be assigned to an unmonitored location; the daily-detection component and its station intercept remained part of the observation model. The existing citywide HEX grid contained 13,714 reporting cells. For each HEX centroid, environmental variables were calculated within a 250-m-radius circle, matching the ecological neighbourhood used to fit the model. HEX polygons were used only to store and display predictions.

The three potential-host models were first projected to every HEX centroid and the 52 weeks of the annual clock. Within each of three host-posterior imputations, every host prediction was standardised using the same full-data mean and standard deviation used for the corresponding Koel fit. The resulting host draws were used in the selected H4 P0 formula without changing its structure. We then combined 200 Koel coefficient draws from each imputation, giving 600 equally weighted posterior predictions for every HEX-week. Weekly acoustic opportunity was summarised only after these draws had been pooled. Calling-density and sampled-minute acoustic-exposure predictions used the selected components defined in S5 without changing their structure; they did not require a new model fit at each HEX.

The completed projection contained 713,128 rows (13,714 HEXes × 52 weeks), with finite ordered posterior quantiles and all probabilities within [0,1]. Supplementary Data 13 summarises the projection requirements and quality checks.

**S6.2 Defining the Prediction Interpretation Domain**

We defined a Prediction Interpretation Domain (PID) to separate locations where a terrestrial or vegetated interpretation was clear from built, bare or land–water mosaics that required caution. PID is a reporting-support label and does not use predicted Koel activity.

Dynamic World class probabilities were averaged within each 250-m circle. The dominant class was the class with the largest mean probability; no minimum probability threshold was imposed. A HEX was classified as PASS when its dominant class was trees, grass, shrub and scrub, flooded vegetation or crops. It was UNCERTAIN when the dominant class was built, bare or water but the circle was not entirely categorical permanent water. A HEX was EXCLUDED when ESA WorldCover identified effectively the whole circle as permanent water. Missing or non-finite required inputs would produce INVALID DATA, although no final-grid HEX had this status.

The final grid contained 5,588 PASS, 7,223 UNCERTAIN and 903 EXCLUDED HEXes. EXCLUDED cells were outside the reporting domain and were treated as missing, not as zero acoustic opportunity or zero exposure. Complete rules are given in Supplementary

Data 13.

#### S6.3 Measuring statistical support for city predictions

Area of Applicability (AOA) asked whether a city HEX resembled the environments represented by the 31 monitoring stations. Because H4 uses predictions from the host models, the comparison included all static environmental inputs required by the host and Koel models: tree cover, built cover, distance to recurrent water, elevation, grass cover, major-road density and tree-edge area. The first four entered the Koel model directly; the remaining variables were required by one or more potential-host models. Host predictions were not added again to the AOA distance.

Each variable was standardised using the 31-station mean and standard deviation. For target HEX $h$, the dissimilarity index was the equal-weight root-mean-square distance to the nearest training station:

$$DI_h = \min_i \left[ \frac{1}{7} \sum_{k=1}^{7} (x_{hk}^* - x_{ik}^*)^2 \right]^{1/2}.$$

We also counted how many predictors fell outside their training-station minimum–maximum ranges. The reference thresholds were obtained by leaving out each station in turn and measuring its distance to the nearest remaining station: $d_{95} = 1.140$ was the 95th percentile and $d_{\max} = 1.468$ was the maximum leave-one-station distance. A HEX was SUPPORTED when no predictor was outside range and $DI \leq d_{95}$; BOUNDARY when no predictor was outside range and $d_{95} < DI \leq d_{\max}$; EXTRAPOLATED when one predictor was outside range and $DI \leq d_{\max}$; and SEVERELY UNSUPPORTED when at least two predictors were outside range or $DI > d_{\max}$.

The H4 AOA contained 3,186 SUPPORTED, 66 BOUNDARY, 2,493 EXTRAPOLATED and 7,969 SEVERELY UNSUPPORTED HEXes. These seven predictors are static, so each HEX retained one AOA label across all 52 weeks; no weekly majority or worst-week aggregation was applied.

**S6.4 Combining ecological and statistical support**

We combined PID and AOA into four mutually exclusive prediction-support classes. Primary Results comprised PASS HEXes with SUPPORTED or BOUNDARY AOA. Moderate Extrapolation comprised reportable HEXes with EXTRAPOLATED AOA, together with UNCERTAIN HEXes that had SUPPORTED or BOUNDARY AOA. Monitoring Gaps comprised PASS or UNCERTAIN HEXes with SEVERELY UNSUPPORTED AOA. Outside the Reporting Domain comprised EXCLUDED HEXes.

The four classes contained 2,712, 3,033, 7,066 and 903 HEXes, respectively. Primary Results and Moderate Extrapolation HEXes formed the planning-facing reporting population used in S7. Monitoring Gaps identified environments where new surveys would be informative; they were not interpreted as low Koel activity. Neither PID, AOA nor the combined class multiplied or otherwise rescaled weekly acoustic opportunity, calling density or sampled-minute acoustic exposure. Supplementary Data 14 gives counts, clipped map areas and the full cross-tabulation, and fig. S7 maps the two support layers and their combined result.

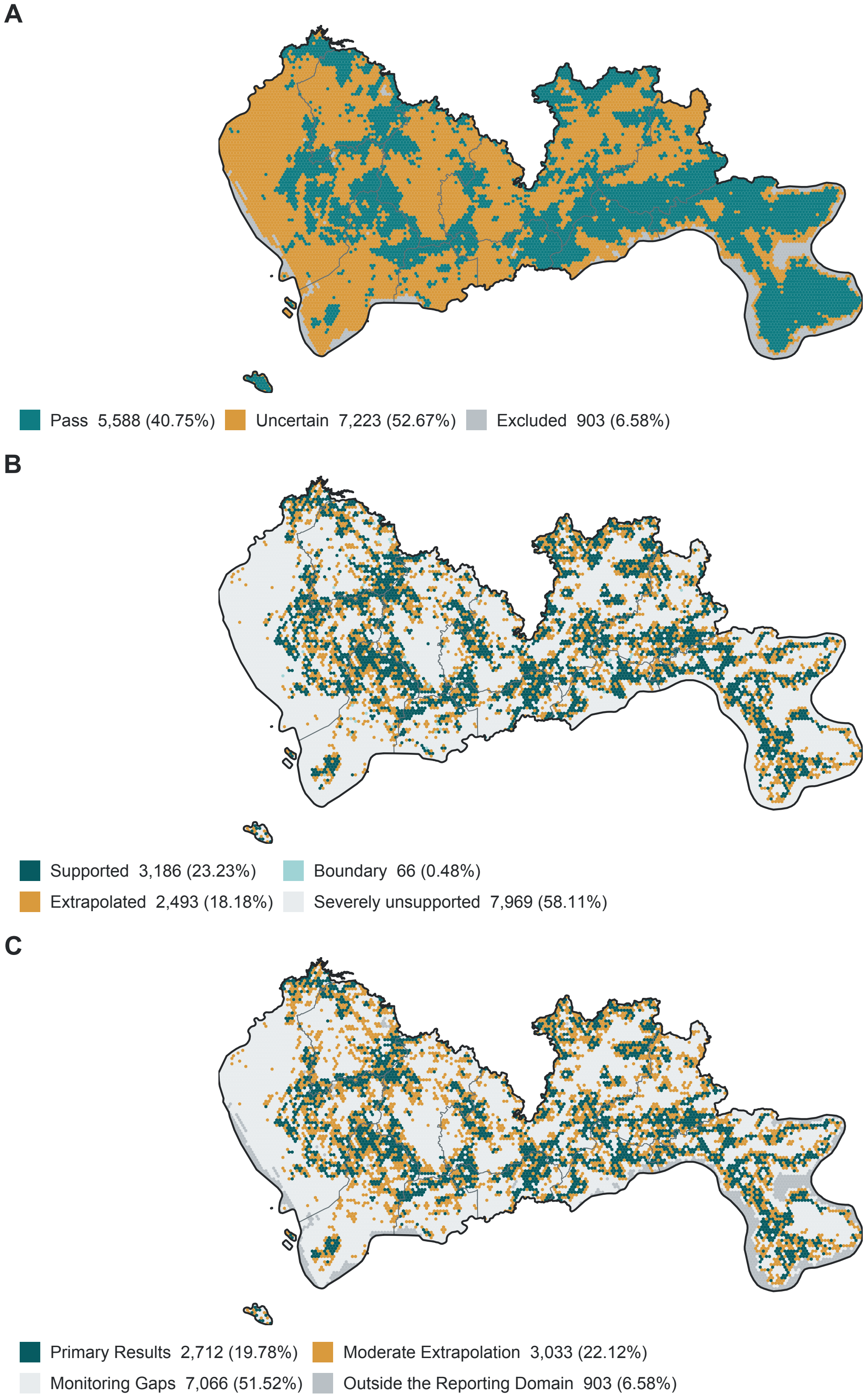
A
Pass 5,588 (40.75%)
Uncertain 7,223 (52.67%)
Excluded 903 (6.58%)
B
Supported 3,186 (23.23%)
Boundary 66 (0.48%)
Extrapolated 2,493 (18.18%)
Severely unsupported 7,969 (58.11%)
C
Primary Results 2,712 (19.78%)
Moderate Extrapolation 3,033 (22.12%)
Monitoring Gaps 7,066 (51.52%)
Outside the Reporting Domain 903 (6.58%)

Fig. S7. Prediction Interpretation Domain and statistical support for citywide three-host predictions. (A) PID separates ecologically interpretable cells, uncertain built/bare/land–water mosaics and categorical permanent-water exclusions. (B) Area of Applicability compares the seven-variable environmental setting of each HEX with the 31 monitoring stations. (C) Their cross-classification yields Primary Results, Moderate Extrapolation, Monitoring Gaps and locations Outside the Reporting Domain. Counts and percentages use all 13,714 HEXes. The same AOA label applies to all 52 weeks because only static environmental variables enter the distance calculation. Support classes accompany predictions but do not multiply or rescale weekly acoustic opportunity, calling density or sampled-minute acoustic exposure. Monitoring Gaps indicate unsupported environments, not low Koel activity; excluded HEXes are outside the reporting domain rather than zero predictions.

### Supplementary Methods S7. Overlap with urban functions and sensitive places

#### S7.1 Restricting summaries to the planning-reporting population

We evaluated urban overlap only in the planning-facing population defined in S6: Primary Results + Moderate Extrapolation. This population contained 5,745 of the 13,714 fixed reporting HEXes. Monitoring Gaps were excluded because their environments were poorly represented by the monitoring network, and permanent-water exclusions remained Outside the Reporting Domain. These filters selected where summaries were reported; they did not alter weekly acoustic opportunity, daily detectability, calling density or sampled-minute acoustic exposure.

All urban layers were linked to the fixed HEX reporting grid described in S6. This was a reporting operation rather than a change in ecological scale. Model predictors still described 250-m-radius circular neighbourhoods around each HEX centroid, whereas urban functions and sensitive places were assigned to the HEXes in which they occurred.

#### S7.2 Mapping sensitive places

Sensitive-place locations were extracted from a dated OpenStreetMap (OSM) PBF snapshot whose latest object edit was 30 March 2026 (© OpenStreetMap contributors; Open Database Licence). We extracted tagged nodes, ways and relations and assigned them to six non-overlapping classes:

1. School and childcare;

2. Higher education;

3. Hospital;

4. Other healthcare and care;

5. Park and garden; and

6. Accommodation.

Polygon features were converted to an interior representative point. Source type, object identifier, version and last-edited timestamp were retained. Exact source duplicates were removed. Named features in the same class with the same normalised name were treated as duplicate map representations when their representative points were within 50 m; 13 such duplicates were removed. Points were clipped to the Shenzhen district union and assigned to the containing reporting HEX. No 250-m buffer was added around a point.

The completed city layer contained 3,653 POIs after duplicate removal. Of these, 763 occurred in Primary Results or Moderate Extrapolation HEXes and entered the numerical summaries. Each retained POI received equal weight. Source tags, category definitions and quality checks are given in Supplementary Data 15 (Table S15A–B).

**S7.3 Mapping residential, commercial and industrial areas**

Residential, commercial and industrial areas were extracted from `landuse=residential`, `landuse=commercial` and `landuse=industrial` polygons in the same dated OSM snapshot. Valid polygons were clipped to the Shenzhen district union and dissolved within each class. We then intersected each class with the Primary Results and Moderate Extrapolation HEXes and calculated the actual polygon–HEX overlap area.

Functional-zone summaries were weighted by this overlap area. The retained overlaps covered 29.23 km² of residential land, 4.61 km² of commercial land and 28.78 km² of industrial land. A HEX could contribute to more than one class when different mapped polygons occurred within it. The overlap operation did not replace the 250-m circular neighbourhood used by the ecological model. Supplementary Data 15 (Tables S15A and S15C) record the source, spatial rule and retained areas.

**S7.4 Summarising acoustic activity within each urban group**

We combined the completed posterior draws from S5 and S6 before calculating group summaries. For posterior draw $s$, week $t$ and urban group $g$, group-weighted weekly opportunity was

$$\bar{\psi}_{gt}^{(s)} = \frac{\sum_h w_{hg}\, \psi_{ht}^{(s)}}{\sum_h w_{hg}},$$

where $w_{hg}$ was the number of POIs of class $g$ remaining in HEX $h$ after duplicate removal, or the polygon-overlap area for a functional zone. POIs therefore contributed equally after deduplication, whereas functional zones contributed in proportion to mapped area.

For each of the six POI classes and three functional zones, we used 100 matched posterior draws from each of the three host imputations, giving 300 draws in total. Weekly-opportunity maps in S6 used 200 draws per imputation. The urban summaries comprised:

1. mean weekly acoustic opportunity across W01–W52;

2. mean weekly acoustic opportunity across W10–W30;

3. the week with the highest posterior-mean opportunity;

4. expected Koel-positive sampled minutes among the 24 scheduled samples from 04:00 to 08:00, averaged across W15–W21; and

5. the posterior probability of at least one positive sampled minute among those same 24 samples.

The two dawn metrics were derived from the fitted beta-binomial process. The dawn window comprised 24 scheduled samples on a reference day. Expected counts and window probabilities were calculated within each week and posterior draw, then averaged across W15–W21 before group summaries were formed. We did not convert a mean minute probability with an independence formula. Values are posterior medians with 95% credible intervals in Supplementary Data 15 (Table S15B-C) and fig. S8.

These summaries describe spatial co-location between mapped urban receptors or functions and modelled Koel acoustic activity under the study sampling design. They do not directly measure human presence, continuous calling duration, sound pressure, perceived loudness, annoyance, exposure dose, habitat use or causal effects of urban land use. OSM coverage also varies among places, so the absence of a mapped POI does not establish the absence of a facility.

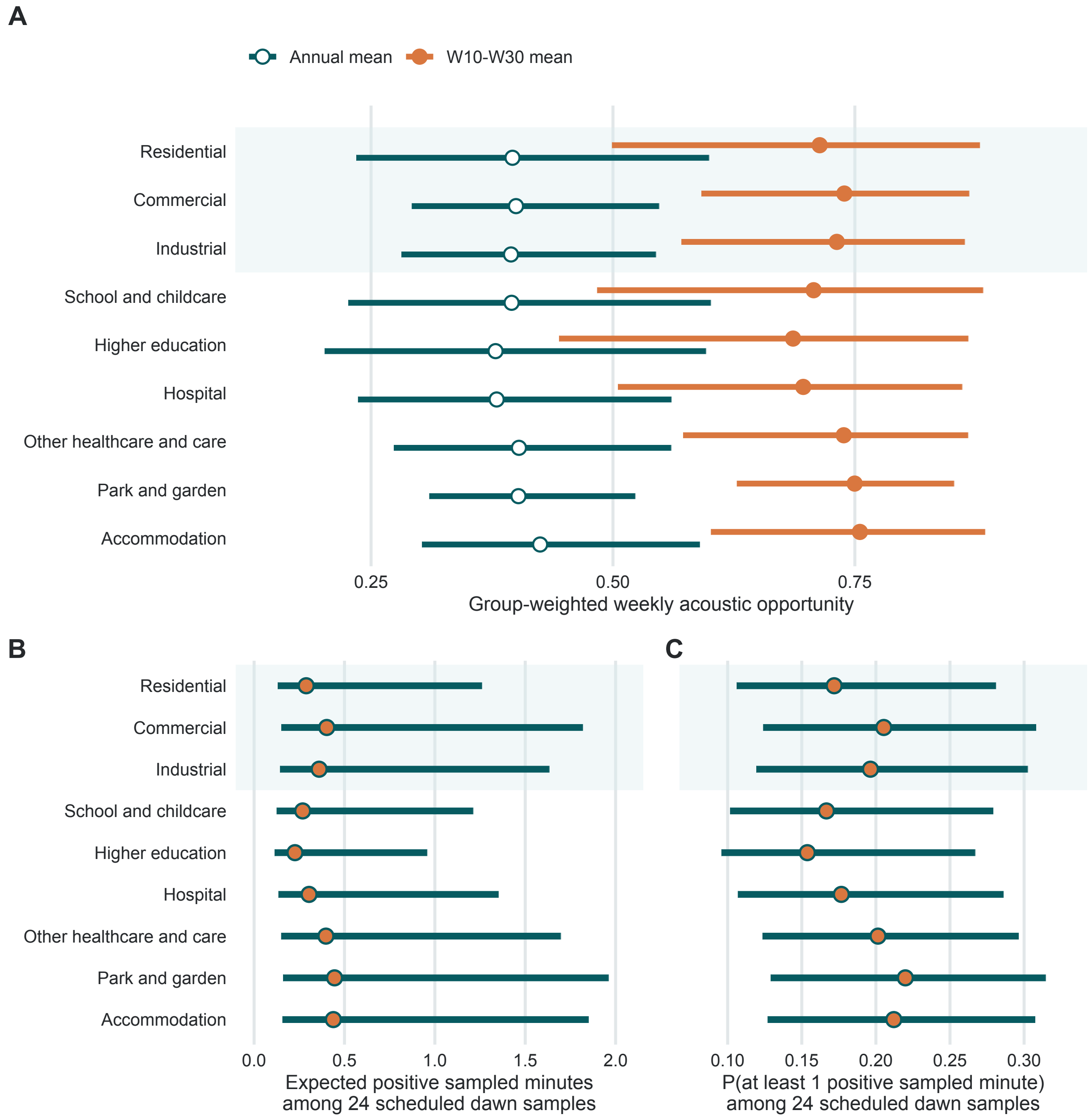

Fig. S8. Modelled Koel acoustic activity overlapping mapped urban functions and sensitive places. Summaries used the 5,745 HEXes classified as Primary Results or Moderate Extrapolation. Urban-function zones were weighted by polygon–HEX overlap area, POIs received equal weight, and uncertainty was propagated across 300 posterior draws. (A) Annual and W10–W30 weekly acoustic opportunity for three functional zones and six sensitive-place classes. (B) Expected Koel-positive sampled minutes among the 24 scheduled samples from 04:00 to 08:00, averaged over W15–W21. (C) Probability of at least one positive sampled minute in the same dawn window, calculated draw by draw from beta-binomial zero-count probabilities with positive-day conditioning. Points are posterior medians and lines are 95% credible intervals. These are modelled co-location summaries under the study sampling design, not direct measures of human exposure, perceived loudness or disturbance. Urban layers: © OpenStreetMap contributors, ODbL.

### Supplementary Methods S8. Designing the next PAM deployment

#### S8.1 Candidate monitoring cells

We used the limitations identified in S6 to define two routes for new passive acoustic monitoring stations. A reporting HEX was eligible when it was inside the Prediction Interpretation Domain (PASS or UNCERTAIN), had complete values for all seven environmental predictors required by the three-host modelling sequence, and did not contain a PAM station. After excluding cells containing existing stations, 7,065 of the 7,066 Monitoring Gap HEXes entered the Coverage Expansion candidate pool; 5,715 Primary Results and Moderate Extrapolation HEXes entered the Prediction Validation candidate pool. We required each new site to be at least 4.627 km from the existing network and previously selected sites. This was the median nearest-neighbour separation among the 31 stations. The resulting pools contained 2,734 and 2,017 candidates, respectively.

#### S8.2 Expanding environmental coverage

The Coverage Expansion route targeted Monitoring Gaps, where ecological interpretation was allowed but the H4 predictor combination was severely unsupported. We temporarily added each candidate to the seven-variable training reference and updated predictor ranges and nearest-neighbour dissimilarity. Its score was the number of gap cells moving below the fixed maximum leave-one-site-out dissimilarity threshold defined in S6. We selected the largest current gain, updated the reference, removed nearby candidates and repeated the calculation. This prevented overlapping one-site gains from being summed. Adding the first candidate to the environmental reference brought 5,123 gap cells prospectively within the support criterion; the first 15 Coverage Expansion candidates covered 6,993 of 7,066 gap HEXes under the same prospective calculation. The illustrative 15-station portfolio used the first nine Coverage Expansion candidates and six Prediction Validation candidates; those nine expansion sites represented 6,884 gap cells (97.4%).

#### S8.3 Validating uncertain predictions

The Prediction Validation route operated within Primary Results + Moderate Extrapolation HEXes. For each candidate, we calculated the mean width of the H4 P0 weekly $\psi$ 95% credible interval over W15-W21. Wider intervals indicated a larger precision gap during the prespecified seasonal peak, not higher Koel activity. We selected the largest remaining width under the same spacing rule. A selected site identifies where new observations could test an uncertain but reportable prediction; it does not predict the resulting interval reduction.

#### S8.4 Planning relevance, deployment size and field checks

Mapped sensitive places and urban functional zones provided planning-sensitive context.

They broke exact ties after the scientific objective, followed by distance from existing stations and HEX identifier. Fig. S9 also shows a public-facing layer combining the stronger within-layer scientific score with mapped planning-sensitive context; it did not determine the sequences.

We evaluated illustrative deployment sizes of 5, 10 and 15 new stations. Proportional allocation by the two spacing-eligible candidate pools gave 3 + 2, 6 + 4 and 9 + 6 Coverage Expansion and Prediction Validation sites, respectively. Portfolios were nested as deployment size increased. This is a sequential heuristic, not a globally optimal network. Every centroid requires field checks for access, permission, power, security, local noise and recorder placement. Candidate rules and route-specific rankings are reported in Supplementary Data 16, and fig. S10 shows the resulting nested portfolios.

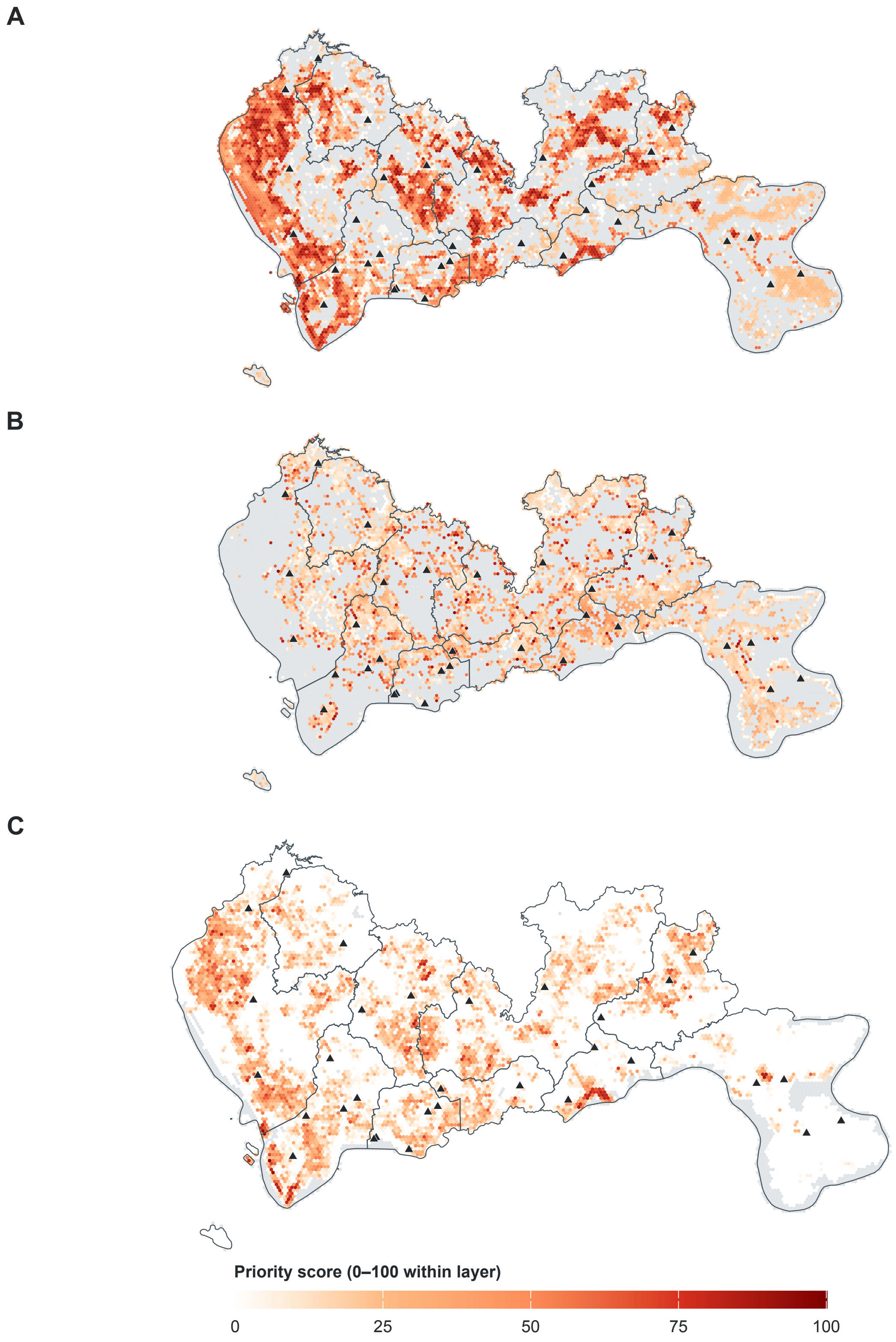


Fig. S9. (A) Coverage Expansion priority, representing prospective gain in H4

environmental applicability within Monitoring Gaps. (B) Prediction Validation priority, representing mean W15–W21 weekly-opportunity interval width within Primary Results and Moderate Extrapolation. (C) A public-facing display layer combining scientific priority with mapped planning-sensitive context; this layer did not determine site selection. Scores are scaled from 0 to 100 within each layer. Grey cells are outside that layer's candidate pool; white eligible cells have a score of zero. Triangles show existing PAM stations.

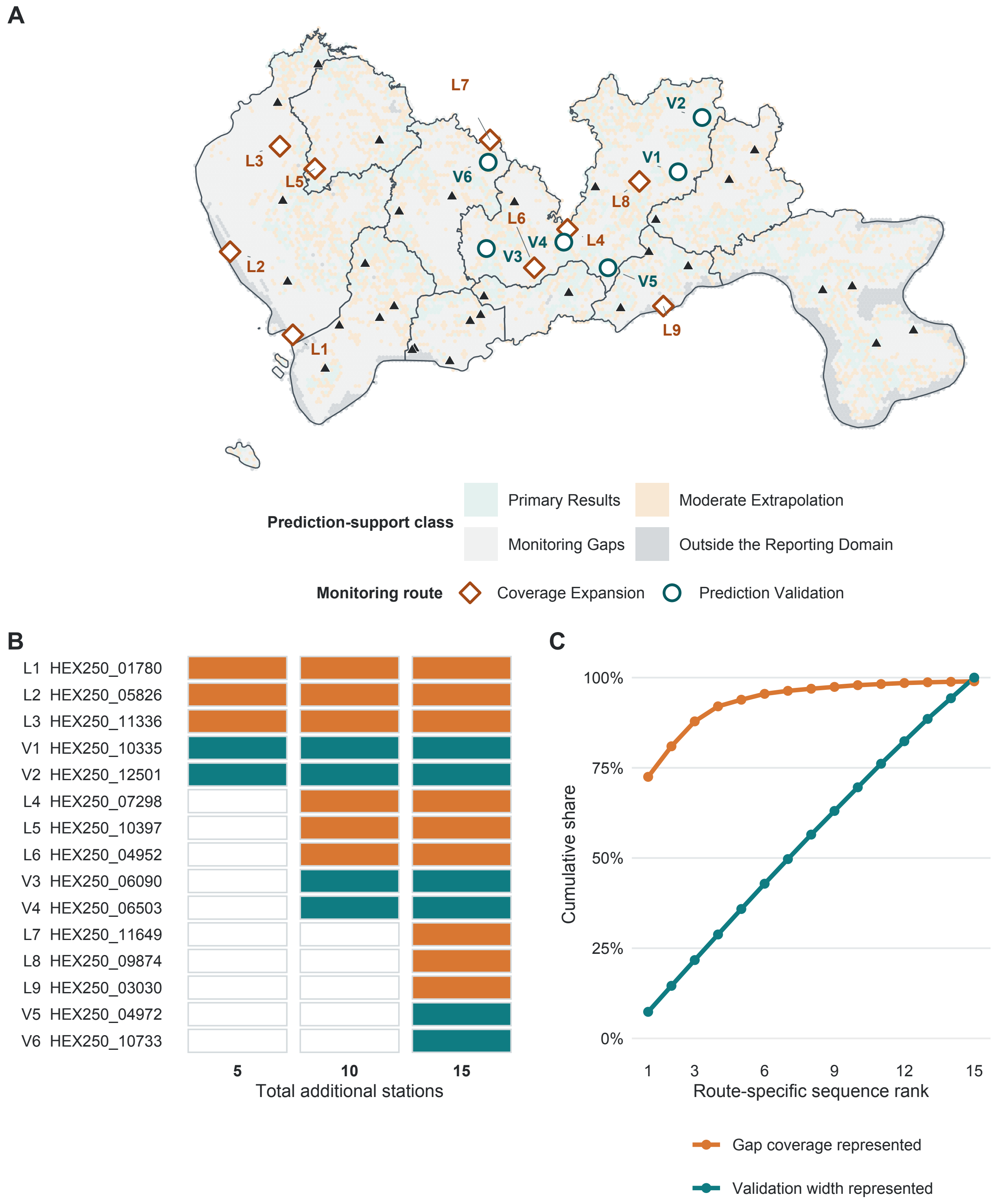


Fig. S10. Sequential candidate sites and stability across deployment sizes. (A) maps the 15-site portfolio, comprising nine Coverage Expansion and six Prediction Validation sites.

(B) shows retention within the nested 5-, 10- and 15-site deployment sizes. (C) shows the cumulative environmental coverage represented by the Coverage Expansion sequence and the cumulative share of the top-15 validation-uncertainty widths represented by the validation sequence. The latter is a prioritisation summary, not an estimate of realised uncertainty reduction. Black triangles show existing PAM stations; orange diamonds and teal circles identify Coverage Expansion and Prediction Validation candidates. Labels give route-specific rank.

### Supplementary Methods S9. Computation and reproducibility

#### S9.1 Software and computing environment

Statistical analyses were run in R 4.5.2 on an arm64 macOS platform. Occupancy models were fitted with `spOccupancy` 0.8.0 (*49*). Custom daily and hourly hurdle models were fitted with `cmdstanr` 0.8.0 and CmdStan 2.38.0 (*48*). Data handling, posterior summaries, model comparison and spatial processing used `data.table` 1.18.2.1, `posterior` 1.6.1, `loo` 2.9.0, `sf` 1.1.0 and `terra` 1.9.1. Raster summaries were calculated in Google Earth Engine, and OpenStreetMap extracts were processed with `osmium-tool` 1.19.0 and projected vector operations in R (*47*). The acoustic-identification architectures and their development data are documented separately in Supplementary Data 3. Supplementary Data 17 (Table S17A) gives the software identities and their analytical roles.

#### S9.2 Reproducible model settings

Each fitted model was saved with its formula, training-fold identifier, random seed, MCMC settings and identifiers of its input files. In spOccupancy fits, occurrence and detection coefficients had Normal priors with mean 0 and variance 2.25 (standard deviation 1.5). In the custom Stan joint models, the corresponding priors had mean 0 and standard deviation 2.25. Formal occupancy fits used four chains. Full-data Koel, Corvid and laughingthrush fits used 30,000 iterations per chain, with 10,000 burn-in iterations and thinning every 20 iterations. The retained Sturnid host refit and whole-station validation fits used 60,000 iterations per chain, with 20,000 burn-in iterations and the same thinning interval. Stan fits used four chains and 750 or 1,000 warm-up and sampling iterations (*53*). Host uncertainty was represented by three independent posterior imputations. Within each validation fold, host fitting, standardisation and Koel fitting used training stations only; complete histories from withheld stations were scored.

Occupancy fits were clean when $\hat{R} \leq 1.05$ and bulk and tail effective sample sizes were at least 400; qualified acceptance required $\hat{R} \leq 1.10$ and both sizes at least 200. Stan fits also required no divergent transitions or maximum-treedepth hits, with $\hat{R} \leq 1.01$ for the formal joint-model fits. Supplementary Data 17 (Table S17B) gives all priors, seeds and draw schedules.

**S9.3 Data sources and reproducibility**

The sources, access dates and licences for spatial datasets are listed in Supplementary Data 7 and 15. Coordinates for all 31 monitoring stations are provided in Supplementary Data 1. Model identifiers and reproducibility checks are summarised in Supplementary Data 17 (Table S17C).
