## Supplementary material for "Urban planning leaves an audible legacy in a megacity’s bird soundscape": Description of Additional Supplementary Files

The following 17 Excel files accompany the manuscript.

Supplementary Data 1. Monitoring stations and survey coverage

Station coordinates, acoustic sampling coverage, Asian Koel detections and assignments to the fixed whole-station cross-validation folds.

Supplementary Data 2. Recording effort and detection-history construction

Reconciliation of recording effort, positive detections, valid non-detections and missing surveys used to construct the daily detection histories.

Supplementary Data 3. Acoustic-identification workflow

Model-development data, shared validation results and filtering rules used for sound-event detection, spectrogram localisation and species identification.

Supplementary Data 4. Daily-detection model comparison

Candidate observation-model definitions and whole-station predictive comparisons evaluating recording effort, acoustic conditions and their temporal structure.

Supplementary Data 5. Daily-detection model checks

Checks of survey inclusion, model design, leverage and sensitivity supporting the retained daily-detection model.

Supplementary Data 6. Model variables and reporting quantities

Definitions, symbols, sources and construction rules for model responses, observation covariates, landscape predictors, potential-host variables and derived reporting quantities.

Supplementary Data 7. Spatial data sources and processing

Spatial-data products, versions, public identifiers and geoprocessing steps used to derive environmental predictors and urban reporting layers.

Supplementary Data 8. Potential-host model development and validation

Species membership, response definitions, candidate models, retained model structures and whole-station validation results for the three potential-host groups.

Supplementary Data 9. Koel acoustic-opportunity model development

Model-development sequence, landscape–season candidate comparisons and the roles of host, coordinate and random-effect structures in association and spatial-transfer analyses.

Supplementary Data 10. Spatial-transfer validation and scale sensitivity

Landscape-neighbourhood comparisons, whole-station predictive metrics, paired model contrasts and the validation and residual-spatial diagnostic protocol.

Supplementary Data 11. Joint acoustic-model components and calling density

Response layers, daily and hourly calling-density candidates, model roles, validation results and posterior checks for the joint Koel acoustic model.

Supplementary Data 12. Acoustic-metric definitions and reference summaries

Definitions, denominators, units and interpretation of model outputs and derived acoustic metrics, together with reference summaries for the retained model.

Supplementary Data 13. Citywide projection and prediction-support rules

Projection inputs and quality checks, Prediction Interpretation Domain and Area of Applicability criteria, and rules for combining them into prediction-support classes.

Supplementary Data 14. Citywide prediction-support summaries

Grid counts and mapped areas of prediction-support classes, cross-tabulation of the two support assessments, and definitions of the classification and reporting populations.

Supplementary Data 15. Acoustic activity in planning-sensitive urban settings

Urban-layer sources and overlay rules, with posterior acoustic-activity summaries for sensitive-place classes and residential, commercial and industrial zones within supported reporting areas.

Supplementary Data 16. Additional monitoring-site selection

Candidate eligibility, prioritisation rules, sequential site lists and nested deployment scenarios for Coverage Expansion and Prediction Validation.

Supplementary Data 17. Software, model settings and reproducibility records

Software names and versions, model priors, sampling settings, random seeds, diagnostic criteria and reproducibility checks.
